# Tumour-derived glucocorticoids amplify ROR1 for targeting with glucocorticoid-resistant ROR1 CAR-T cells

**DOI:** 10.1101/2025.01.28.632923

**Authors:** Marc P. Schauer, Justus Weber, Peter Spieler, Barbara Altieri, Michael Rade, Leon Gehrke, Stefan Kircher, Silviu Sbiera, Max Kurlbaum, Matthias Kroiss, Katja Kiseljak-Vassiliades, Margaret E. Wierman, Thomas Nerreter, Kristin Reiche, Sophia Danhof, Hermann Einsele, Martin Fassnacht, Laura-Sophie Landwehr, Michael Hudecek

## Abstract

Exogenous glucocorticoids (GCs) suppress T cell-based immunotherapy, yet the consequences of endogenous GCs produced or regenerated by tumours remain poorly understood. Here, we show that tumour-derived GCs couple immune evasion to therapeutic antigenicity. In adrenocortical carcinoma, autonomous GC production activates hGR-STAT3 signaling to increase the surface antigen ROR1. This creates an immunological paradox in which GC excess promotes therapeutic antigen expression while suppressing T cell-mediated antitumour activity. Selective hGR-knockout renders ROR1 CAR-T cells GC-resistant while preserving GC-driven ROR1 expression in tumour cells, resulting in durable tumour control *in vivo*. Non-endocrine pancreatic and triple-negative breast cancers recreate this circuit through HSD11B1-mediated GC recycling, which is further induced by CAR-T cell-derived cytokines under immune pressure. Thus, GC-resistant CAR-T cells exploit a tumour-derived endocrine program that couples immune suppression to therapeutic antigenicity.

**Significance:** Our study reveals that endogenous autocrine glucocorticoid signaling couples immune suppression to therapeutic antigen amplification, creating a tumour-intrinsic vulnerability that can be exploited by glucocorticoid-resistant CAR-T cells.

## INTRODUCTION

Exogenous glucocorticoids (GCs) – administered therapeutically to manage inflammation, chemotherapy-related toxicity, or immune-related adverse events^1,2^ – suppress T cell viability, effector function, and persistence through glucocorticoid receptor (GR)-mediated transcriptional reprogramming.^3,4^ Their negative impact on the efficacy of adoptive T cell therapies and immune checkpoint inhibitors has been documented across multiple clinical and preclinical contexts.^5–7^ What has received substantially less attention is whether endogenous GCs – produced autonomously by tumour cells rather than exogenously administered – exert the same capacity to undermine T cell-based immunotherapy, and whether they do so through mechanisms that extend beyond paracrine immunosuppression.

GCs confer their immunosuppressive effects through GR-mediated interference with pro-inflammatory transcription factors and transactivation of immunosuppressive target genes, broadly reshaping the activation, differentiation, and function of both adaptive and innate immune cells.^3,8,9^ Tumour-derived GCs differ from their exogenous counterparts in two immunologically important aspects: (1) they are constitutively present throughout the anti-tumour immune response, locally enriched within the tumour microenvironment, and scale dynamically with tumour burden rather than with a physician’s dosing decision while (2) creating an immunosuppressive niche that is autonomous and a barrier to therapeutic efficacy.

Endogenous, tumour-derived GC production is not a rare phenomenon.^10,11^ In adrenocortical carcinoma (ACC) – the prototypic steroidogenic malignancy – the majority of cases present with excessive autonomous de novo GC production that leads to increased intra-tumoural and systemic GC concentrations.^12^ In a previous study, we have shown that GC excess in ACC affects the phenotype and function of infiltrating immune cells, and is linked to the failure of immunotherapy with tumour infiltrating lymphocytes (TIL) and immune checkpoint inhibitors.^13,14^ Elevated intra-tumoural GC concentrations are not restricted to endocrine tumours and have also been reported in non-endocrine cancers including TNBC, melanoma, and PDAC,^15,16^ in which GCs accumulate through recycling from inactive metabolites in tumour cells that express HSD11B1, as well as in cancer-associated fibroblasts and tumour-associated macrophages.^17–19^ A largely uncharted area of investigation relates to the effect of these endogenous GCs on chimeric antigen receptor (CAR-)T cell therapy. Unlike TIL- and T cell receptor (TCR-)based approaches that depend on peptide-MHC complexes for target recognition, CAR-T cells recognize surface antigens independently of MHC and are thus not subject to GC-mediated suppression of antigen presentation.^20^ What has not been considered is whether the same autocrine GC signaling that can suppress MHC expression on tumour cells may alter surface antigens in a pro-tumourigenic context, potentially amplifying CAR-T cell target density while remaining invisible to conventional T cell surveillance. If tumour-derived GCs simultaneously suppress the immune response and reshape the antigen landscape of cancer cells, the net consequence for CAR-T cell therapy would depend critically on the balance between these two effects and on whether engineering GC resistance into the CAR-T cell product itself could resolve this tension.

We addressed these questions in ACC as the paradigmatic model of endogenous GC excess in cancer, with GC production that is cell-autonomous, quantifiable, and directly linked to immune exclusion. We show that tumour-derived GCs fuel an autocrine signaling loop that drives overexpression of the oncogenic surface antigen receptor tyrosine kinase-like orphan receptor 1 (ROR1) – examined here as a representative example of a GC-regulated tumour antigen, with the broader implication that hormonal regulation of surface antigen expression may be operative across hormonally active cancers. Importantly, this biology is not restricted to endocrine cancers with autonomous GC production. We show that non-endocrine cancers, including TNBC and PDAC, recreate the same GC-hGR/STAT3-ROR1 axis through HSD11B1-mediated local GC recycling. The same GC excess that amplifies ROR1 simultaneously suppresses infiltrating T cells through paracrine signaling, creating an immunological paradox in which autocrine signaling and its consequences for antigen amplification and immune evasion are mechanistically coupled. We resolve this paradox by generating GC-resistant ROR1 CAR-T cells through CRISPR/Cas9-mediated human GR (hGR) knockout, leaving autocrine GC signaling and its antigen-amplifying consequences unaffected in tumour cells while constitutively protecting CAR-T cells from paracrine immunosuppression. ^hGR-KO^ROR1 CAR-T cells were effective against ACC, TNBC and PDAC, whereas GC excess diminished the viability and function of non-gene-edited ROR1 CAR-T cells. This design converts tumoural GC release from a therapeutic liability into a precision opportunity and shows that endogenous tumour-derived GC excess is not simply an immunological barrier but a targetable biological axis with direct therapeutic consequences for the design of next-generation cellular immunotherapies.

## RESULTS

### ROR1 is highly expressed in ACC with GC excess

We analyzed RNA transcriptome array data to identify genes that are overexpressed in ACC compared to normal adrenal glands (nAG) and that encode surface antigens.^21^ This analysis identified *ROR1* as a top candidate that was markedly overexpressed in ACC (**Fig. 1a**). We confirmed overexpression of *ROR1* transcripts in clinically annotated primary ACC samples **(Table 1)**, even though we also noted a range with high and low expressers **(Fig. 1b and Extended Data Fig. 1a)**. In multivariate analyses, high *ROR1* expression in primary ACC was associated with higher ACC aggressiveness based on the ENSAT clinical staging system, Ki67 proliferation index and the histopathological Weiss score. Furthermore, its expression correlated with an increased risk for rapid recurrence after first-line therapy **(Fig. 1c-f and Extended Data Fig. 1b)**.^21–24^ We assessed clinical ACC samples by immunohistochemistry and found that ROR1 expression was stable between diagnosis and relapse, and similar between primary and metastatic ACC lesions **(Fig. 1G, Suppl. Table S1).** Among the patients with the highest ROR1 expression as determined by immunohistochemistry, all patients had clinically presented with systemic GC excess. Therefore, we grouped samples according to the presence vs. absence of GC excess (ACC^GC+^ vs. ACC^GC-^) and found a clear segregation of the two groups, with higher ROR1 expression in ACC^GC+^ (**Fig. 1h,i)**. Of note, in the subgroup of ACC patients with androgen rather than GC excess, ROR1 expression was similar to ACC^GC–^.

**Fig. 1.**
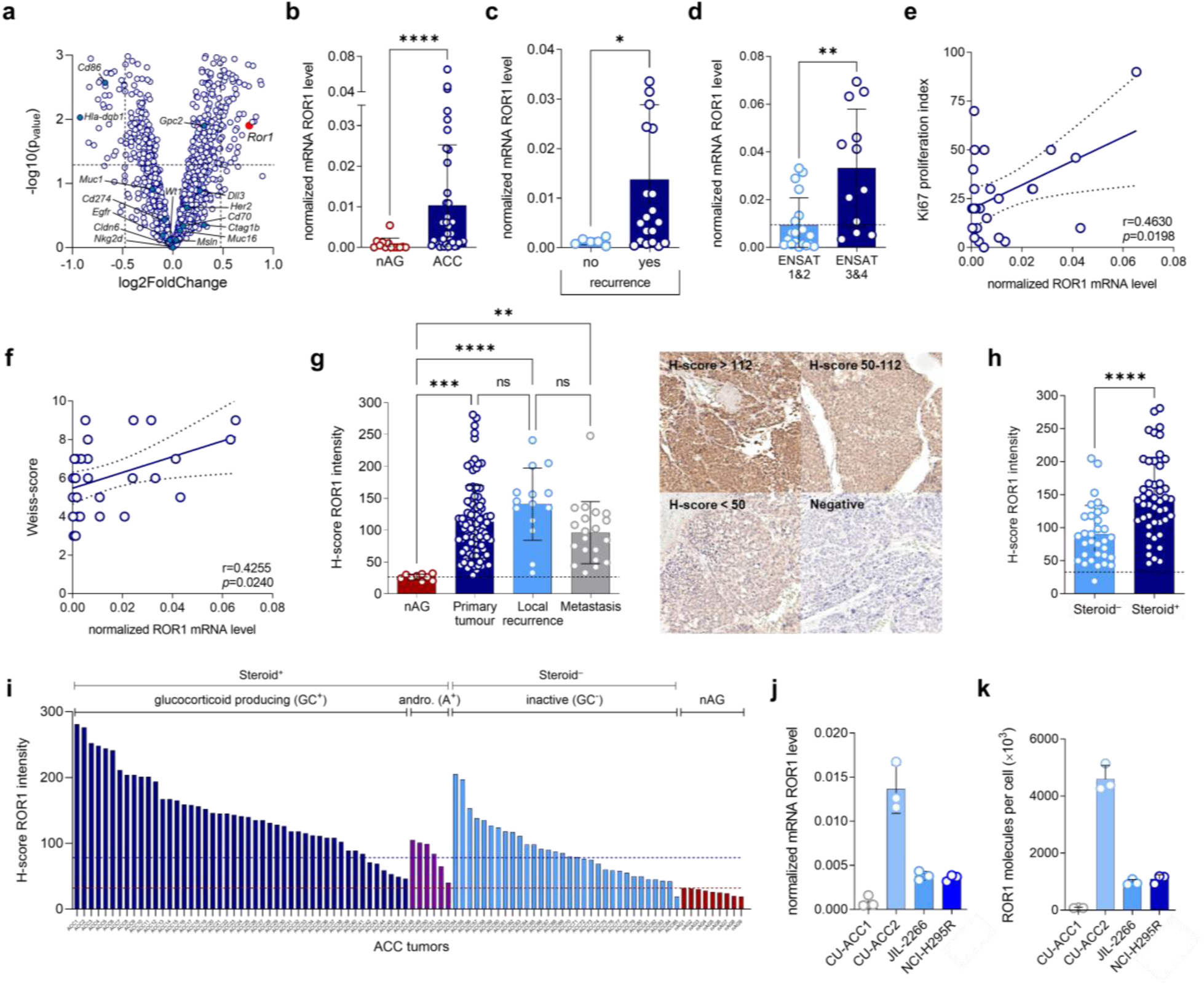
ROR1 is highly expressed in ACC with GC excess. (**a**) Volcano plot of mRNA gene expression assessed by RNA sequencing analysis comparing normal adrenal glands (nAG) with adrenocortical carcinoma (ACC) samples. ROR1 gene expression and other known CAR/TCR targets are annotated. The horizontal line is at an adjusted *p* value of 0.05. (**b**) Comparison of ROR1 mRNA expression (normalized to α-tubulin) in normal adrenal glands (nAG) compared to ACC samples assessed by quantitative real-time PCR (qRT-PCR). (**c**) Comparison of ROR1 mRNA expression in ACC patients with and without disease recurrence. (**d**) Association between ROR1 mRNA expression in ACC patients with low (ENSAT 1&2) and high (ENSAT 3&4) ENSAT tumour stage. (**e**) Association of ROR1 mRNA expression with Ki67 proliferation index and (**f**) histopathological Weiss-score. (**g**) Immunohistochemical ROR1 protein analysis comparing H-scores of nAG, primary ACC tumours, local recurrences and metastases. Representative pictures are shown alongside. (**h**) ROR1 protein expression comparing steroid hormone inactive (Steroid^−^) with steroid hormone producing (Steroid^+^) ACC tumours. (**i**) Waterfall plot comparing ROR1 protein expression in ACC samples with glucocorticoid (GC^+^) excess, androgen excess (andro^+^), inactive (GC^−^) ACC tumours and nAGs. Mean value of ROR1 expression in inactive ACC tumours (blue line) and the highest ROR1 expression in nAGs (red line) are annotated. (**j**) ROR1 mRNA expression of the four human ACC cell lines assessed by quantitative real-time PCR (qRT-PCR) (**k**) ROR1 molecules per cell (×10^3^) of all four ACC cell lines assessed by quantitative flow cytometry (n=3). Statistical analyses were performed using unpaired *t*-test (two-tailed) (**b,c,d,h**), Pearson correlation (**e,f**), repeated measures one-way ANOVA with Dunnett’s test for multiple comparisons (**g**) and were considered significant if *p* value was \**p*<0.05,\*\**p*<0.01,\*\*\**p*<0.001, \*\*\*\**p*<0.0001; ns, not significant.

We confirmed ROR1 expression in four ACC cell lines - CU-ACC1 & CU-ACC2^25^, NCI-H295R^26^ and JIL-2266^27^ - that we used in addition to clinical ACC samples in our subsequent experiments **(Fig. 1j,k and Extended Data Fig. 1c,d)**. Each of these ACC cell lines secreted a distinct steroid hormone profile (**Extended Data Fig. 1e**). We quantified *ROR1* transcripts by qRT-PCR and found a similar range of *ROR1* expression between our patient cohort and the set of ACC cell lines (**Extended Data Fig. 1a**). Further quantitative analyses by RNAscope and by flow cytometry confirmed ROR1 RNA and protein expression in each of the ACC cell lines (**Fig. 1k and Extended Data Fig. 1c**). We employed super-resolution *d*STORM microscopy to analyze the distribution and density of ROR1 protein on the surface of ACC cell lines and determined CU-ACC2 to be ROR1^high^ (0.428 clusters/µm^2^), JIL-2266 (0.116 clusters/µm^2^) and NCI-H295R (0.239 clusters/µm^2^) to be ROR1^medium^, and CU-ACC1 to be ROR1^low^ (0.054 clusters/µm^2^), using ROR1^+^ (Jeko-1) and ROR1^-^ (K562) hematologic cancer cell lines as reference (**Extended Data Fig. 1d,f-i**). Taken together, our analyses identified ROR1 as a highly expressed cancer antigen and marker of aggressive phenotype in ACC. ROR1 expression was particularly high in the subgroup of patients with GC excess (ACC^GC+^).

### GC inhibitors diminish ROR1 expression in ACC

We sought to probe the efficacy of ROR1-specific CAR-T cells against ACC and reasoned that the *de novo* production of immunosuppressive GCs from ACC cells would compromise their anti-tumour function **(Extended Data Fig. 2a)**. Therefore, we treated ACC cells with various GC inhibitors comprising inhibitors of steroidogenesis (i.e. mitotane, metyrapone or ketoconazole) and a hGR inhibitor (relacorilant) prior to treatment with ROR1 CAR-T cells. Unexpectedly, we found a decrease in cytolytic activity and cytokine release from ROR1 CAR-T cells against GC inhibitor treated vs. non-treated ACC cells **(Fig. 2a and Extended Data Fig. 2b-d)**. Upon further analyses, we found decreased expression of *ROR1* transcripts and of ROR1 cell surface protein after GC inhibitor treatment in ACC cells **(Fig. 2b and Extended Data Fig. 2e, 3a-c)**. Mitotane, the most commonly used GC inhibitor in patients with ACC, exerted the most potent effect and conferred a >90% decrease in *ROR1* transcripts and converted almost all ACC cells from being ROR1^+^ to ROR1^-^ as assessed by flow cytometry and *d*STORM super-resolution microscopy **(Fig. 2b and Extended Data Fig. 3a-c)**. The effect of mitotane was dose-dependent, persistent but reversible **(Fig. 2c)**. After discontinuation of mitotane treatment, *ROR1* transcripts recovered to baseline and subsequently, ROR1 protein expression on the surface of ACC cells also returned to baseline **(Fig. 2d)**. When dexamethasone was added to the culture medium, the time to recovery of *ROR1* transcripts and ROR1 protein to baseline was substantially shorter, and even exceeded baseline expression **(Fig. 2d,e)**. A similar effect was confirmed for metyrapone, ketoconazole and relacorilant, respectively **(Fig. 2b and Extended Data Fig. 3a-c)**. These data suggested that GCs were involved in regulating ROR1 expression in ACC. To corroborate our data, we turned to our collection of annotated clinical ACC samples and indeed, found significantly lower *ROR1* transcript and ROR1 protein expression in samples obtained from ACC patients that had received mitotane or other GC inhibitors prior to sampling **(Fig. 2f**, **Table 1)**. The decrease in ROR1 expression was particularly evident between matched samples from primary and metastatic ACC lesions that were obtained prior to and after GC inhibitor therapy, respectively **(Fig. 2g,h and Extended Data Fig. 3d)**. Taken together, these data show that treatment with GC inhibitors diminishes ROR1 expression in ACC^GC+^ and support a causal relationship between GC excess and ROR1 overexpression in ACC.

**Fig. 2.**
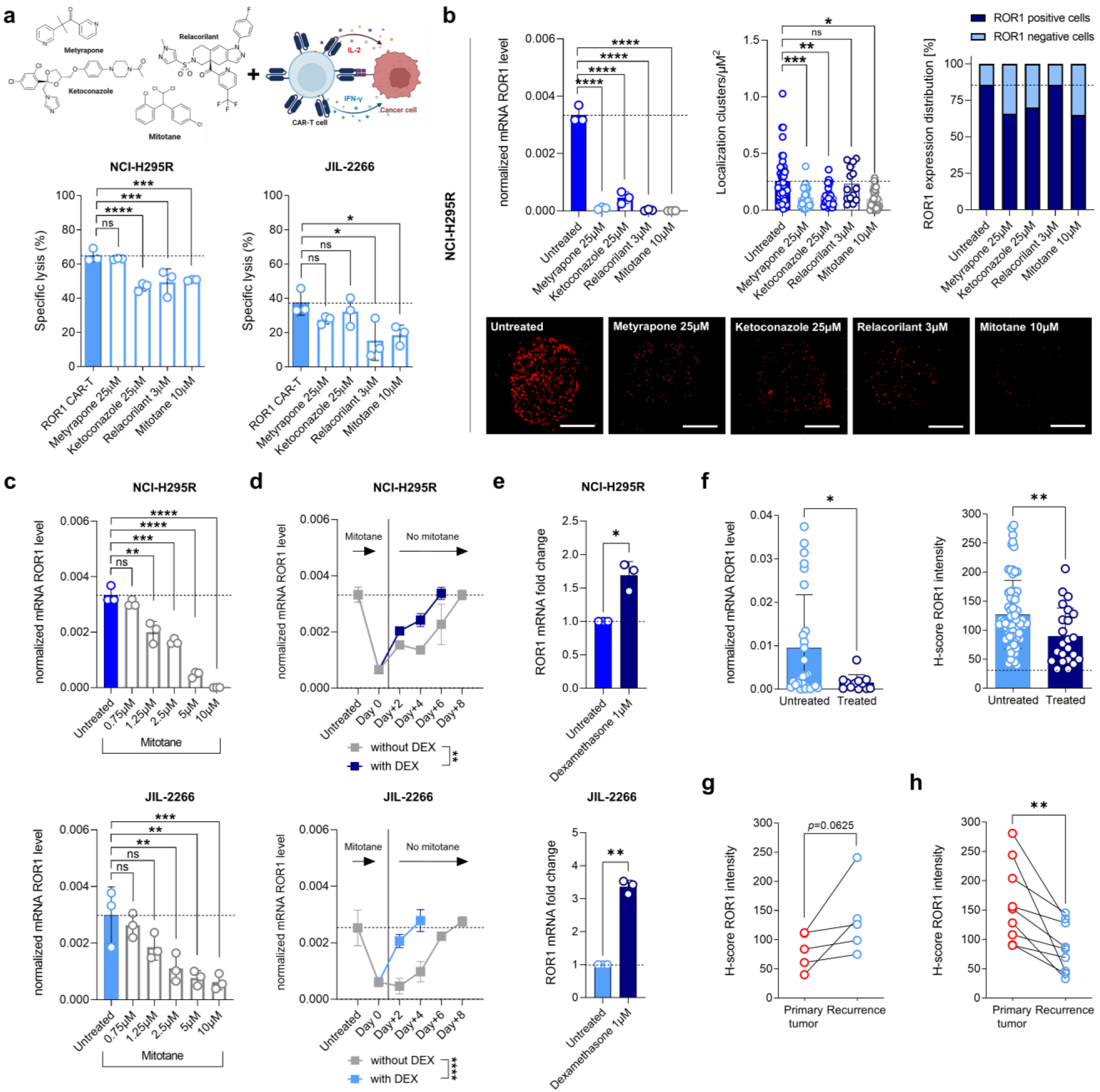
GC inhibitors diminish ROR1 expression in ACC. (**a**) Treatment scheme used for the in vitro experiment. Specific lysis of ROR1 CAR-T cells targeting the steroidogenic NCI-H295R and non-steroidogenic JIL-2266 ACC cell line in 3D cell culture after 48 hours of co-culture with glucocorticoid (GC) inhibitors. Specific lysis of CAR-T cells in combination with the individual drugs in presence was normalized for cell lysis with the individual drugs without CAR-T cells and compared to lysis of UTD T cells, n=3). (**b**) ROR1 downregulation on NCI-H295R cells after 48 hours of co-culture with pharmaceutical inhibitors of GC effector functions. *ROR1* mRNA expression was assessed using qRT-PCR (n=3) and surface density (clusters/µm^2^) and percentage of ROR1^+^ cells were assessed by single molecule high resolution microscopy (*d*STORM, n=30). Representative pictures are shown alongside. (**c**) ROR1 mRNA expression after treatment with mitotane for 48 hours with different dose-dependencies (n=3). (**d**) ROR1 mRNA expression after abolishing exposure to mitotane at different time points (n=3). (**e**) ROR1 mRNA expression of NCI-H295R cells after 48 hours of co-culture with 1µM of dexamethasone (n=3). (**f**) ROR1 mRNA and protein expression of ACC patientś tissue samples comparing untreated samples with those that received GC inhibitors prior to surgical removal of the relapsed tumour lesion. ROR1 protein expression in matched ACC tumour samples before (**g**) and after **h**, receiving GC inhibitors. Statistical analyses were performed using repeated measures one-way ANOVA with Dunnett’s test for multiple comparisons (**a,b,c**), unpaired t-test (two-tailed) (**f**), paired t-test (**e,g,h**), and were considered significant if *p* value was \**p*<0.05,\*\**p*<0.01,\*\*\**p*<0.001, \*\*\*\**p*<0.0001; ns, not significant; hGR, human glucocorticoid receptor.

### *ROR1* transcription in ACC is regulated by hGR and STAT3

In the next set of experiments, we focused our analyses on ACC cells that we had treated with GC inhibitors to derive further insights into the regulation of ROR1 expression. Treatment with GC inhibitors increased hGR and STAT3 expression at both the transcript and protein level, compared to non-treated ACC cells **(Extended Data Fig. 4a-d, 5a)**. Furthermore, we detected increased amounts of the ROR1 ligand WNT5A, suggesting that a positive feedback loop was active in ACC cells to sustain ROR1 signaling **(Extended Data Fig. 4a-d)**.

The concurrent upregulation of hGR and STAT3 under conditions of GC inhibition, combined with prior reports of hGR/STAT3 transcriptional cooperation in breast cancer^28^ prompted us to investigate whether GC-activated hGR and STAT3 physically interact and jointly regulate ROR1 expression in the context of GC excess, and that pharmacological GC inhibition dissolves this complex, thereby relieving ROR1 transcription (**Extended Data Fig. 4e**). We performed proximity ligation analyses to demonstrate the direct interaction of hGR and STAT3 in ACC cells, and found that treatment with GC inhibitors decreased the amount of hGR/STAT3 complexes **(Fig 3a)**, and decreased ROR1 surface expression **(Fig. 3c)**. We used ACC samples from our clinical cohort to confirm these data and to demonstrate that hGR/STAT3 complexes were significantly more abundant in ACC^GC+^ compared to ACC^GC–^. The data showed that therapy with GC inhibitors in patients abrogated the formation of hGR/STAT3 complexes and expression of ROR1 in both ACC^GC+^ and ACC^GC-^ **(Fig. 3b and Extended Data Fig. 5b).** In all clinical ACC samples and cell lines, the number of hGR/STAT3 complexes strongly correlated with ROR1 expression (**Fig. 3i,j and Extended Data Fig. 6a,b**). To provide definitive evidence for the dependence of ROR1 expression on hGR signaling, we knocked down *hGR* in ACC cells and found a significant reduction in ROR1 expression, whereas WNT5A and STAT3 expression was strongly upregulated **(Fig. 3e,f and Extended Data Fig. 5c,d)**.

**Fig. 3.**
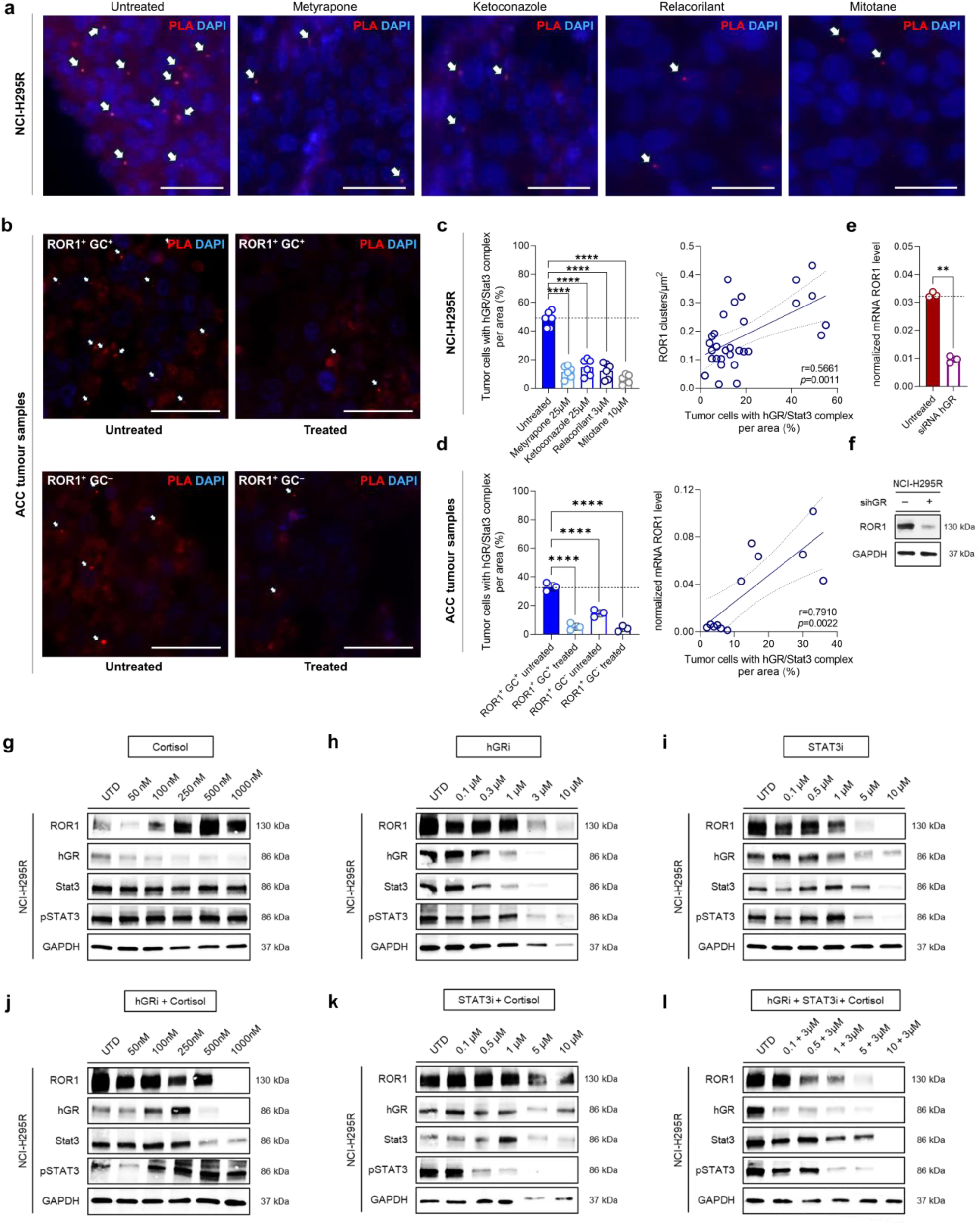
ROR1 transcription in ACC is regulated by hGR and STAT3. (**a**) Representative pictures of Duolink proximity ligation analysis (PLA) assessing hGR/Stat3 protein complex (PLA in red and DAPI in blue) in positive NCI-H295R cells in 3D cell culture before (untreated) and after 48 hours of treatment with different glucocorticoid (GC) inhibitors and in (**b**) primary ACC patient samples with and without GC excess before and after GC inhibitor treatment. Comparison of hGR/Stat3 complex positive tumour cells (per area (%) before and after treatment in (**c**) NCI-H295R and (**d**) primary ACC cells as well as the association between hGR/Stat3 complexes and ROR1 expression. (**e** and **f**) ROR1 expression before and after hGR gene silencing in NCI-H295R ACC cells. (**g-l**) ROR1, hGR, STAT3 and pSTAT3 protein expression in NCI-H295R cells after 48 hours of treatment with either hGR inhibition (relacorilant), STAT3 inhibition (WT1066) and cortisol alone or in combination. Statistical analyses were performed using repeated measures one-way ANOVA with Dunnett’s test for multiple comparisons (**c,d**), Pearson correlation (**c,d**), paired t-test (**j**) and were considered significant if *p* value was \**p*<0.05,\*\**p*<0.01,\*\*\**p*<0.001, \*\*\*\**p*<0.0001; ns, not significant; hGR, human glucocorticoid receptor.

Collectively, these data show a mechanistically consistent link between autocrine GC signaling, hGR/STAT3 complex formation, and ROR1 overexpression in ACC.

### ROR1 upregulation requires synergistic hGR/STAT3 activity

To determine whether hGR and STAT3 each contribute independently to ROR1 transcription or act exclusively through their physical interaction as a composite complex, we treated ACC cells with the hGR antagonist relacorilant, the STAT3 inhibitor WP1066, or their combination, with or without exogenous cortisol (**Fig. 3g-l**). hGR inhibitor dose titration with relacorilant alone reduced ROR1 protein levels by approximately 60% (**Fig. 3h**). Selective STAT3 inhibition alone produced a ROR1 reduction of almost 70% (**Fig. 3i**) confirming that hGR and STAT3 contribute in a co-dependent, non-redundant manner to ROR1 expression. To determine whether cortisol could restore ROR1 expression under these conditions, we added exogenous cortisol to each inhibitor condition. In the presence of hGR inhibition alone, cortisol did not restore ROR1 expression, consistent with the requirement for intact hGR signaling (**Fig. 3j**). Conversely, when STAT3 was inhibited while hGR remained active, cortisol partially restored ROR1 protein levels, suggesting that hGR signaling alone retains limited capacity to sustain ROR1 expression in the absence of STAT3 (**Fig. 3k**). Under combined hGR and STAT3 blockade, cortisol failed to restore ROR1 expression regardless of dose (**Fig. 3l**).

Taken together, these data establish that hGR and STAT3 are jointly and non-redundantly required for GC-driven ROR1 expression – neither factor alone is sufficient, and both must be simultaneously active for full ROR1 expression in ACC.

### ROR1 CAR-T cells with hGR knockout are resistant to ACC-derived GCs

We reasoned that due to the high density of cell surface ROR1, ACC cells ought to be susceptible to recognition and elimination by CAR-T cells directed against the ROR1 antigen, provided that these were sufficiently protected from the immunosuppressive effect of ACC-derived GCs. Therefore, we performed gene-editing to knockout the *hGR* (hGR-KO) and generated GC-resistant ROR1 CAR-T cells (^hGR-KO^ROR1 CAR-T cells; **Fig. 4a-d and Extended Data Fig. 7a,b**). ^hGR-KO^ROR1 CAR-T cells and non-edited ROR1 CAR-T cells had a similar composition with naïve, effector and memory subpopulations at the end of manufacturing (**Fig. 4e**). We performed a GC challenge with a pulse of 100 nM dexamethasone for 72 hours and found a marked drop in T cell viability in non-edited ROR1 CAR-T cells, whereas ^hGR-KO^ROR1 CAR-T cells maintained viability and yield **(Fig. 4d)**. After the dexamethasone pulse, we found that genes involved in T cell activation and signaling, antigen presentation, immune checkpoint pathways and epigenetic modulation were differentially expressed between hGR-KO and non-edited ROR1 CAR-T cells (**Fig. 4f**). Exemplary genes like *FOXO1* and *LEF1* that are responsible for promoting memory function and for preventing exhaustion, respectively, were significantly higher expressed in ^hGR-KO^ROR1 CAR-T cells as compared to non-edited ROR1 CAR-T cells (**Extended Data Fig. 7c,d**).

**Fig. 4.**
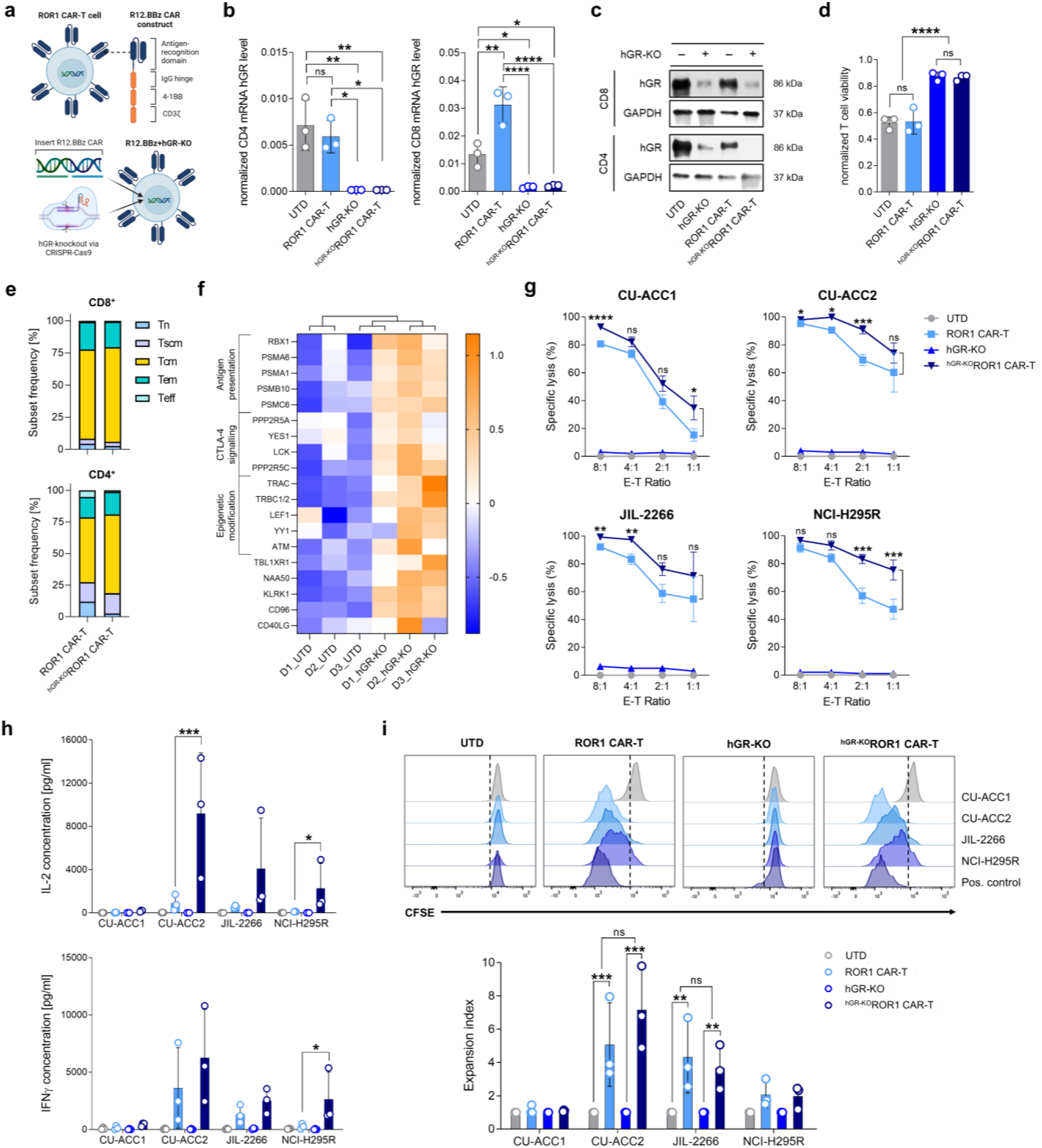
ROR1 CAR-T cells with hGR knockout are resistant to ACC-derived GCs. (**a**) Schematic design of ROR1 CAR-T cells (R12.BBz) with human glucocorticoid receptor knockout (hGR-KO) (ROR1 CAR construct and R12.BBz+hGR-KO). Untransduced T cells (UTD) have been used for normalization. (**b**) hGR mRNA (qRT-PCR) and (**c**) protein expression (Western Blot) before and after CRISPR/Cas9-mediated hGR knockout. (**d**) T cell viability after 100 nM dexamethasone treatment for 48 hours. (**e**) ROR1 CAR-T cell subset analysis before and after CRISPR/Cas9-mediated glucocorticoid receptor knockout. (**f**) Heatmap of dysregulated genes in T cells with and without hGR-KO. (**g**) Specific lysis of four ROR1^+^ ACC cell lines with different effector-to-target ratios (E-T) after 24 hours. (**h**) IL-2 and IFN-γ cytokine secretion of ROR1 CAR-T cells with and without hGR knockout after antigen contact for 24 hours was assessed by ELISA in the supernatant. (**i**) Proliferation and expansion of ROR1 CAR-T cells upon antigen contact in all four ACC cell lines after 72 hours assessed by flow cytometry. Statistical analyses were performed using repeated measures one-way ANOVA with Dunnett’s test for multiple comparisons (**b,e**), unpaired *t*-test (two-tailed) (**d**), paired *t*-test (**g,h,i**) and were considered significant if *p*-value was \**p*<0.05,\*\**p*<0.01,\*\*\**p*<0.001, \*\*\*\**p*<0.0001; ns, not significant; ELISA, enzyme-linked immunosorbent assay.

We performed reference testing against the non-steroidogenic lymphoma and leukemia cell lines JeKo1 and K562 and found that ^hGR-KO^ROR1 CAR-T cells exerted similarly potent effector and helper functions as non-edited ROR1 CAR-T cells **(Extended Data Fig. 8a)**. We then tested ^hGR-KO^ROR1 CAR-T cells against each of the steroidogenic ACC cell lines and recorded superior cytolytic activity compared with non-edited ROR1 CAR-T cells, particularly evident against NCI-H295R cells that produce the highest amount of GCs (**Fig. 4g and Extended Data Fig. 1e**). ^hGR-KO^ROR1 CAR-T cells also maintained their ability to produce and release IFN-γ and IL-2, and to undergo productive proliferation after antigen-specific stimulation **(Fig. 4h,i and Extended Data Fig. 8b,c).** Importantly, ^hGR-KO^ROR1 CAR-T cell activation and function remained firmly dependent on the presence of and correlated with the amount of ROR1 antigen on ACC cells **(Extended Data Fig. 9a,b)**.

### GC-resistant ROR1 CAR-T cells induce durable remission of ACC^GC+^ *in vivo*

We sought to translate our findings into an *in vivo* system and established ACC tumours in immunodeficient mice through subcutaneous inoculation of steroidogenic NCI-H295R cells. We confirmed that ACC tumours were ROR1^+^, were in a highly proliferative state by Ki67 staining, were fully developed as evidenced by CD31 expression on tumour vasculature, and expressed several characteristic steroidogenic enzymes and transcription factors (**Fig. 5a and Extended Data Fig. 10**). Serum analyses confirmed GC excess, which correlated with ACC tumour volume (r=0.6571, *p*=0.0016) (**Fig. 5b**). On day 14 after tumour inoculation, mice were randomized to treatment groups to receive a single intravenous dose of 1×10^6^ modified vs. non-modified T cells **(Fig. 5c)**. All of the mice that were treated with ^hGR-KO^ROR1 CAR-T cells showed a marked reduction in tumour burden (overall response rate 100%) within seven days of T cell infusion and remained in complete remission for 5 consecutive weeks (**Fig. 5d-f**). In the group of mice treated with non-edited ROR1 CAR-T cells, 2/7 mice that experienced tumour regression, which was stable for up to 2 weeks. However, at 3 weeks after T cell infusion, all of the mice presented with progressing tumours **(Fig. 5d-f)**. In the group of mice treated with ^hGR-KO^T cells (non-CAR-modified) or untransduced (UTD) T cells, all of the mice showed continuous tumour progression **(Fig. 5d-f)**. With further follow-up, 6/7 mice treated with ^hGR-KO^ROR1 CAR-T cells remained in either partial or complete remission. There was a significant survival benefit with median overall survival not reached *vs.* 36 days for mice treated with hGR-KO vs. non-edited ROR1 CAR-T cells, respectively (*p*<0.0001) **(Fig. 5d,e)**. ^hGR-KO^ROR1 CAR-T cells showed consistent engraftment, *in vivo* proliferation and persistence **(Fig. 5g-l)**, as well as efficient migration into ACC lesions **(Fig. 5k,l)**. There was long-term persistence of ^hGR-KO^ROR1 CAR-T cells **(Fig. 5h)** with a high frequency of stem central memory (T_scm_) and central memory (T_cm_) T cells, a low frequency of PD-1/TIM-3/LAG-3 triple-positive T cells, and elevated expression of the activation markers CD25 and CD69 (**Fig. 5i,j**).

**Fig. 5.**
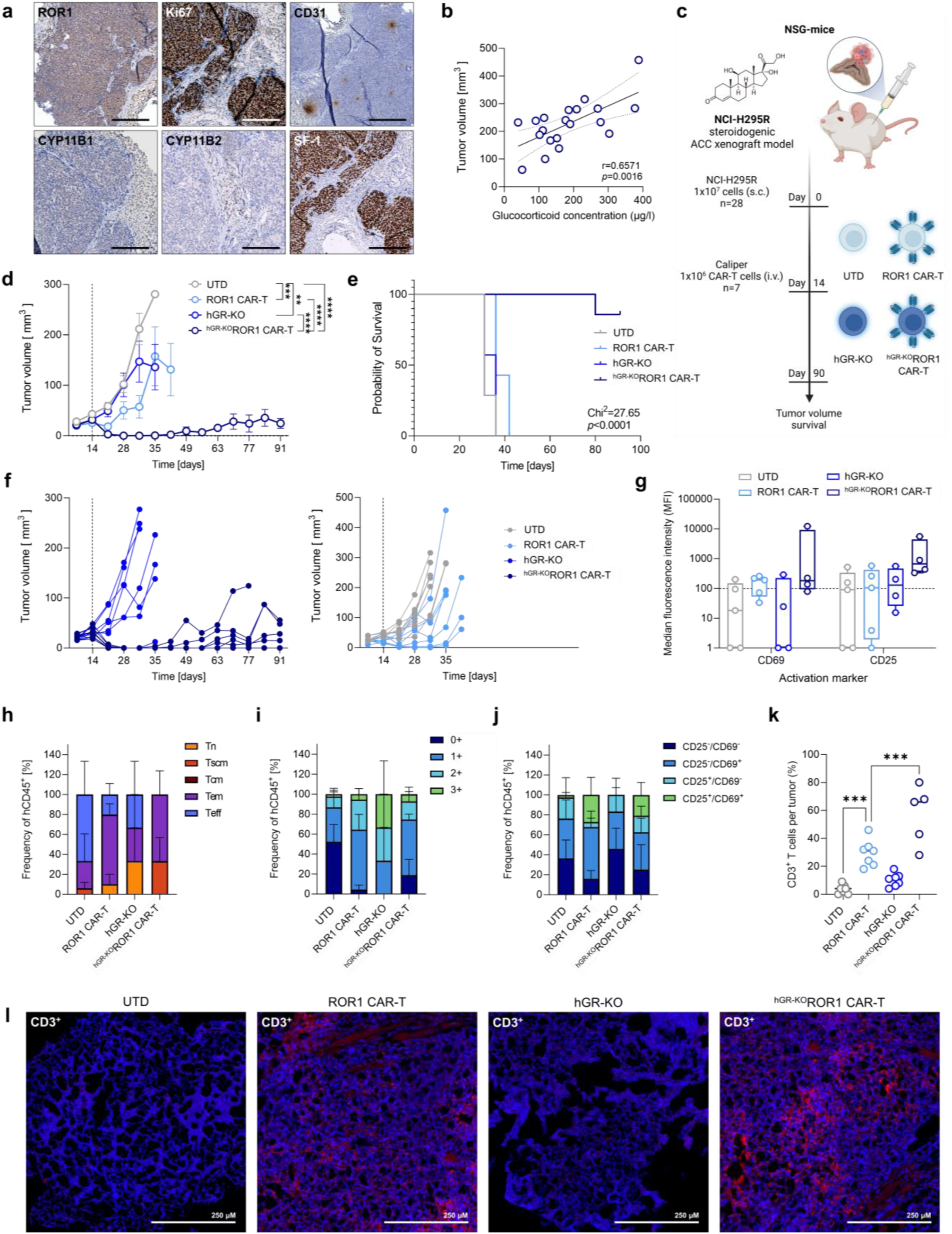
GC-resistant ROR1 CAR-T cells induce durable remission of ACC^GC+^ *in vivo*. (**a**) Characterization of ACC xenografts for ACC specific origin and protein markers. (**b**) Association between glucocorticoid serum levels (Corticosterone) as determined by LC-MS/MS and tumour volume of all mice. (**c**) Treatment scheme used for the *in vivo* experiment. NSG mice bearing steroidogenic NCI-H295R ACC xenografts were treated intravenously on day 14 after tumour inoculation with 1×10^6^ CAR-T cells. (**d**) Quantification of tumour burden for each treatment group measured by digital caliper measuring (n=7 mice per group). (**e**) Survival curve for all tumour-bearing mice treated with different CAR-T/T cell modifications. (**f**) Quantification of tumour burden for each individual mouse and in their different treatment groups measured by digital caliper measuring (n=7 mice per group). (**g**) Median fluorescence intensity of activated and persistent CAR-T cells that are positive for CD25 and CD69 positive for tEGFR transduction marker. (**h**) T cell subset phenotypes at all different endpoints from all four treatment groups. (**i**) Numbers and frequencies of CAR-T cells that have been positive for canonically expressed exhaustion markers. (**j**) Numbers and frequencies of activated CAR-T cells that are positive for CD25 and CD69 at their different endpoints. (**k**) Immune infiltration of CD3^+^ T cells from all different treatment groups assessed by immunofluorescence. Representative pictures for each group are shown in (**l**) alongside. Statistical analyses were performed using Pearson correlation (**b**), repeated measures one-way ANOVA with Dunnett’s test for multiple comparisons (**g-j**), mixed-effects analysis (**d,f**), Log-rank (Mantel-Cox) test (**e**), unpaired *t*-test (**k**) and were considered significant if *p* value was \**p*<0.05,\*\**p*<0.01,\*\*\**p*<0.001, \*\*\*\**p*<0.0001; ns, not significant; hGR, human glucocorticoid receptor; LC-MS/MS, liquid-chromatography-mass-spectrometry.

Collectively, the data show that ^hGR-KO^ROR1 CAR-T cells maintain their function in the presence of immunosuppressive GCs in ACC and outperform conventional, non-gene-edited ROR1 CAR-T cells *in vitro* and *in vivo*. The data obtained in the setting of ACC provide an example of autocrine GC signaling that regulates ROR1 expression in an endocrine tumour with endogenous, *de novo* production of GCs, while ROR1 CAR-T cells were shielded from the paracrine effects of ACC-derived GCs.

### GC recycling by HSD11B1 augments ROR1 expression in PDAC and TNBC

Non-endocrine cancers can release GCs by recycling hormonally inactive cortisone into hormonally active cortisol through the enzyme HSD11B1 **(Fig. 6a)**.^11,16^ To determine which cell types within the TME drive this recycling activity, we analyzed publicly available single-cell RNA sequencing datasets from healthy and tumoural tissues **(Fig. 6b)**. HSD11B1 expression was selectively enriched in malignant cells, cancer-associated fibroblasts, and tumour-associated macrophages, while T cells and other immune effector populations showed no detectable HSD11B1 expression **(Fig. 6b)**. Analysis of TCGA genomic data further revealed that breast cancer exhibits the highest frequency of HSD11B1 copy-number gains among all analyzed tumour entities, present in 69.65% of cases – compared to 66.04% in hepatocellular carcinoma and 63.32% in lung adenocarcinoma – suggesting that HSD11B1 activity is not only driven by tumoural HSD11B1 upregulation but also genomic amplification **(Fig. 6c)**.

**Fig. 6.**
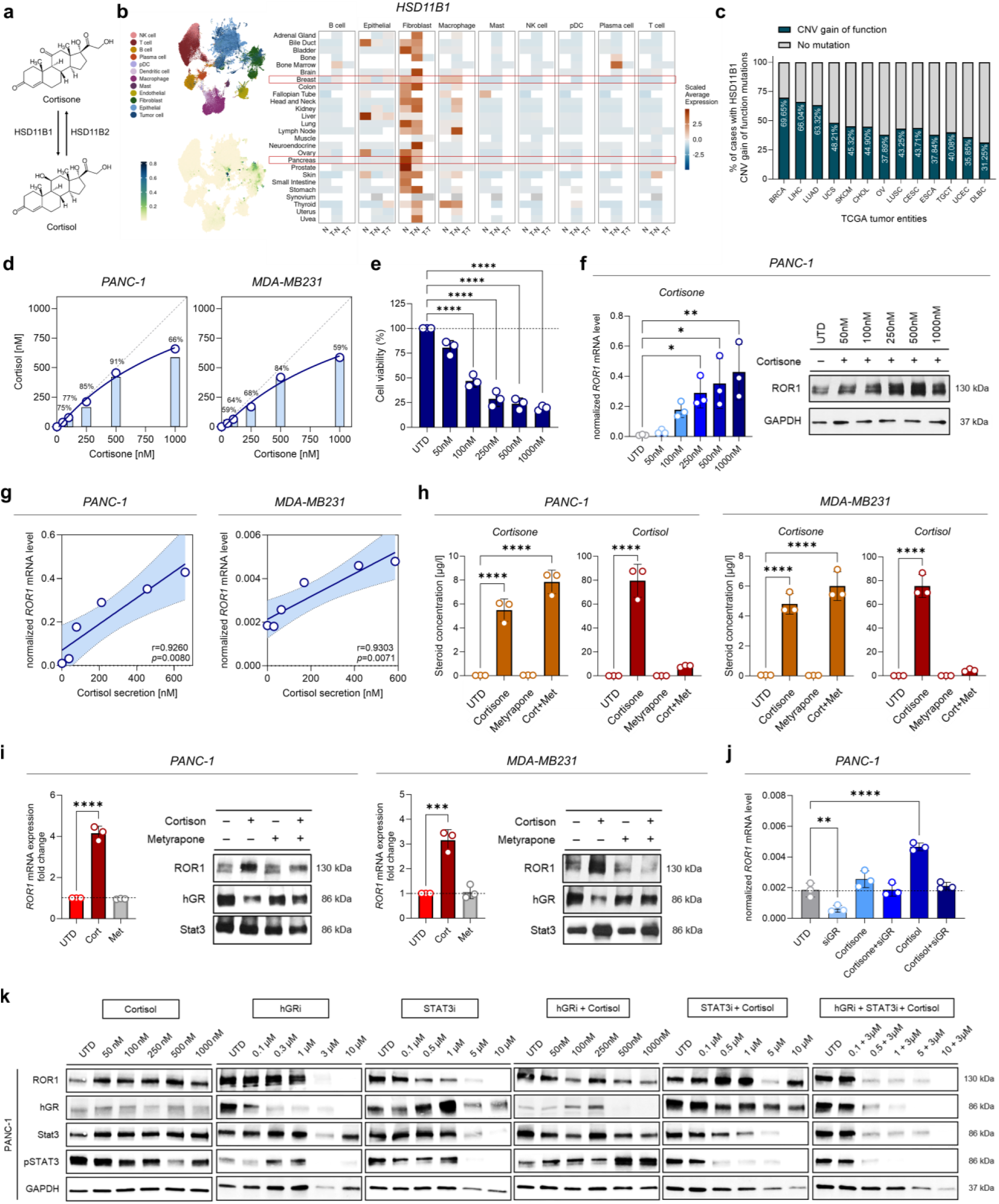
GC recycling by HSD11B1 augments ROR1 expression in PDAC and TNBC. **(a)** Scheme that demonstrates the molecular recouperation from cortisone into cortisol through HSD11B1. **(b)** Single cell RNA-sequencing analyses of HSD11B1 expression derived from a publicly available dataset covering 30 different cancer types across 104 datasets. Pan-cancer tumour-normal landscape on meta-cell resolution including 5 million cells to 300,000 meta-cells (grouping of similar cells (base on the transcriptome profiles)). Meta cells were summarized to hexabins. Color shows average expression of these bins; N: Normal Tissue; T-N: Tumour tissue, Normal cells; T-T: Tumour tissue, Tumour cells. Grey tiles: <10 meta-cells expressing the gene (>0 counts) across all cell types or the average expression is zero across all cell types, White tiles: No data available (**c**) Percentage of HSD11B1 copy-number-gain of function mutations in different solid tumour types extracted from TCGA transcriptomic datasets on cBioPortal. (**d**) PANC-1 and MDA-MB231 induced conversion from cortisone into cortisol as measured by LC-MS/MS after 24 hours of co-culture. (**e**) CAR-T cell viability after treatment with converted tumour cell culture supernatants as shown in Fig. 6d. (**f**) ROR1 mRNA (qRT-PCR) and protein expression (Western Blot) after different doses of cortisone (UTD, untreated control). (**g**) Association between converted cortisol secretion levels and ROR1 mRNA expression. (**h**) Steroid concentrations of cortisone and cortisol with and without adding metyrapone. (**i**) ROR1 mRNA and protein expression of ROR1, hGR and Stat3 after cortisone and metyrapone treatment. (**j**) ROR1 mRNA expression after cortisone and cortisol treatment with and without previous silencing of hGR. (**k**) ROR1, hGR, STAT3 and pSTAT3 proetin expression in PANC-1 cells after 48 hours of treatment with either hGR inhibition (relacorilant), STAT3 inhibition (WT1066) and cortisol alone or in combination. Statistical analyses were performed using one-way ANOVA with Dunnett’s test for multiple comparisons (**e,f,h,I,j**), and Pearson-correlation (**g**) and were considered significant if *p*-value was \**p*<0.05,\*\**p*<0.01,\*\*\**p*<0.001, \*\*\*\**p*<0.0001; ns, not significant.

Given that ROR1 is frequently expressed in both TNBC and PDAC,^29,30^ and that malignant cells represent a major source of intra-tumoural HSD11B1 activity, we reasoned that the autocrine GC signaling loop driving ROR1 overexpression in ACC may be similarly operative in these non-endocrine entities through GC recycling, and that GC-resistant ROR1 CAR-T cells may therefore confer enhanced antitumour reactivity against TNBC and PDAC. First, we treated PANC-1 cells (PDAC) and MDA-MB231 cells (TNBC) with increasing amounts of “inactive” cortisone. Within 24 hours, we could detect “active” cortisol in the culture medium **(Fig. 6b)**. The amount of cortisol had an almost linear correlation to the amount of cortisone that was added to PANC-1 cells and MDA-MB231 cells. The highest amount of cortisol was detected in PANC-1 cells, suggesting a higher cortisone recycling capacity compared with MDA-MB231 cells **(Fig. 6d)**. Without addition of cortisone, there was no cortisol release neither from PANC-1 cells nor MDA-MB231 cells, showing that cortisol was the product of cortisone recycling, and not of *de novo* production **(Fig. 6d).** When we maintained ROR1 CAR-T cells in conditioned medium from cortisone-treated PANC-1 cells and MDA-MB231 cells, respectively, we noted a decrease in T cell viability, supporting that recycled cortisol was hormonally active and immunosuppressive **(Fig. 6e)**. In both PANC-1 cells and MDA-MB231 cells, we found increased expression of *ROR1* transcripts and ROR1 protein following cortisone treatment and cortisone-to-cortisol conversion **(Fig. 6f and Extended Data Fig. 11a,b,c).** *ROR1* expression in PANC-1 cells and MDA-MB231 cells strongly correlated with the amount of cortisol that was recycled and released from these cells **(Fig. 6g)**. To demonstrate the dependence of GC recycling in PANC-1 cells and MDA-MB231 cells on the enzyme HSD11B1, we used metyrapone, a CYP11B1 inhibitor that also inhibits HSD11B1.^31^ Metyrapone treatment efficiently blocked the cortisone-to-cortisol conversion in PANC-1 cells and in MDA-MB231 cells, and prevented the increase in *ROR1* transcript and ROR1 protein expression subsequent to cortisone treatment **(Fig. 6h,i)**. Lastly, when we silenced hGR expression in PANC-1 cells to interfere with autocrine signaling, we observed no upregulation of ROR1 following cortisone and cortisol treatment **(Fig. 6j)**. To confirm that the hGR/STAT3 requirement identified in ACC extends to GC recycling-driven ROR1 regulation in PDAC, we performed rescue experiments in PANC-1 cells. Selective hGR or STAT3 inhibition each reduced ROR1 protein expression by 75%; combined blockade suppressed ROR1 by >90%, and cortisol failed to rescue ROR1 expression when both pathways were simultaneously inhibited **(Fig. 6k)**.

Taken together, these data show that HSD11B1 is catalytically active in PANC-1 cells and in MDA-MB231 cells and fuels the autocrine signaling loop that augments ROR1 expression through GC recycling.

### GC-resistant ROR1 CAR-T cells are effective against PDAC and TNBC *in vivo*

To evaluate *in vivo* efficacy against non-endocrine GC-recycling tumours, we established PANC-1 and MDA-MB231 xenografts in immunodeficient NSG mice. Thereby we eliminated the confounding of an active tumour microenvironment providing a clean experimental system in which all detected GC recycling, ROR1 regulation, and paracrine immunosuppression can be traced directly to the steroidogenic activity of the human tumour cells.

Prior to *in vivo* evaluation, we treated PANC-1 cells and MDA-MB231 cells with cortisone to induce cortisone-to-cortisol conversion and upregulation of ROR1 for targeting with ROR1 CAR-T cells. Prior to functional testing, we maintained ^hGR-KO^ROR1 CAR-T cells and non-edited ROR1 CAR-T cells in PANC-1 vs. MDA-MB231-conditioned medium after cortisone-to-cortisol conversion. ^hGR-KO^ROR1 CAR-T cells were resistant to the recycled GCs and preserved their functional capacity to confer superior effector functions including cytolysis, cytokine production compared with non-edited ROR1 CAR-T cells **(Fig. 7a,b)**. **E**ncouraged by these results, we assessed *in vivo* efficacy in xenograft models of PDAC (PANC-1) and TNBC (MDA-MB231), respectively **(Fig. 7c)**. We confirmed that HSD11B1 converted murine 11-deoxycorticosterone (DOC) into active corticosterone, and also confirmed the presence of DOC in serum of NSG mice **(Extended Data Fig. 11d-f)**. Following tumour engraftment, mice were randomized to treatment groups to receive ^hGR-KO^ROR1 CAR-T cells, non-edited ROR1 CAR-T cells, or UTD T cells, respectively. In both, the PDAC and the TNBC model, ^hGR-KO^ROR1 CAR-T cells conferred superior anti-tumour efficacy compared with non-edited ROR1 CAR-T cells, resulting in a statistically significant survival benefit compared with non-edited ROR1 CAR-T cells and UTD T cells **(Fig. 7d,e)**.

**Fig. 7.**
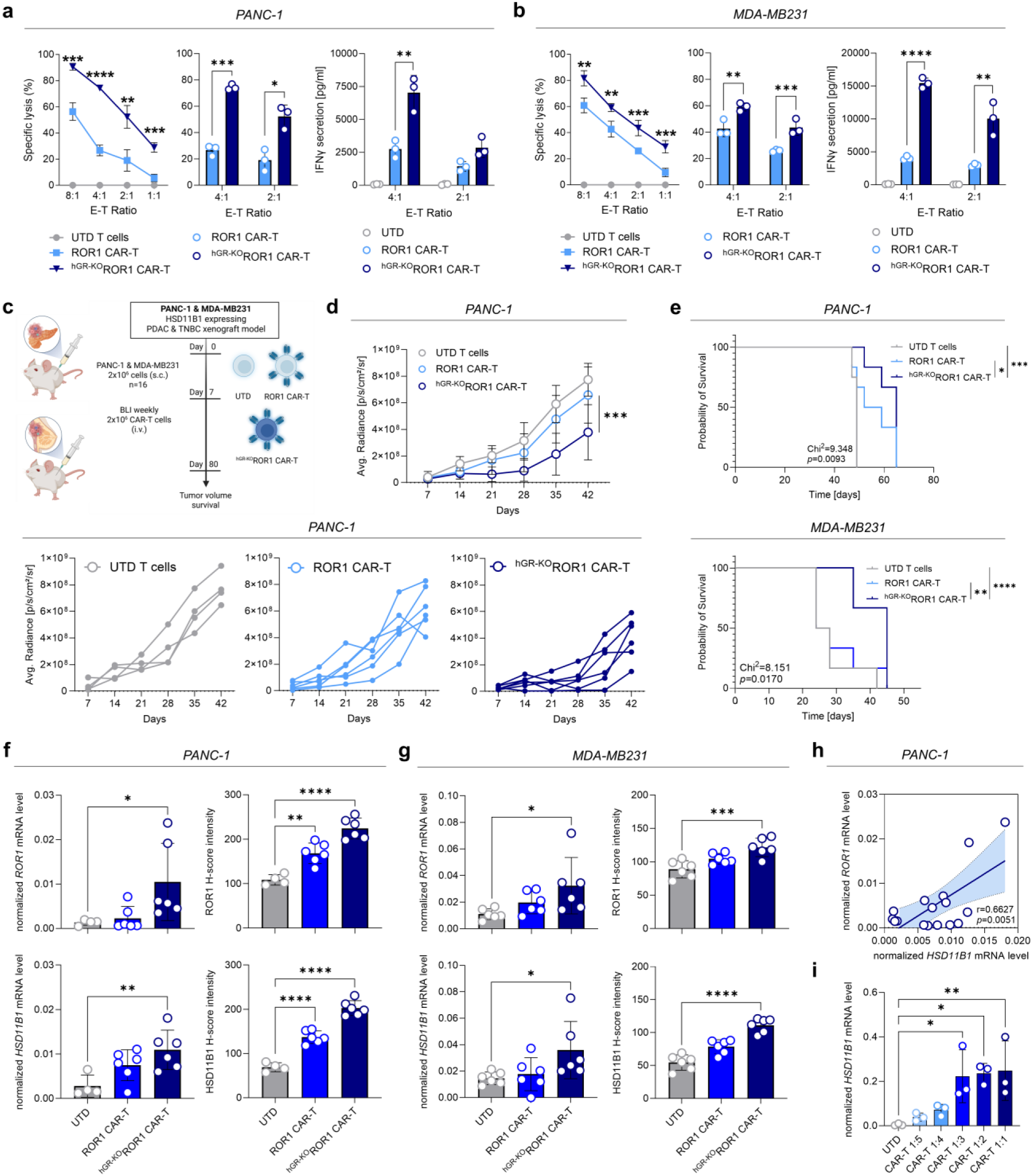
GC-resistant ROR1 CAR-T cells are effective against non-endocrine cancers *in vivo*. (**a**) Specific lysis of ROR1^+^ PANC-1 cells and in (**b**) MDA-MB231 cells with different effector-to-target ratios (E-T) after 24 hours after 24h tumour cell pre-treatment with 500nM of cortisone as well as IFN-γ cytokine secretion of ROR1 CAR-T cells with and without hGR knockout after antigen contact for 24 hours was assessed by ELISA in the supernatant. (**c**) Treatment scheme used for the *in vivo* experiment. NSG mice bearing HSD11B1 expressing PANC-1 (PDAC) and MDA-MB231 (TNBC) xenografts were treated intravenously on day 7 after tumour inoculation with 2×10^6^ CAR-T cells. (**d**) Quantification of tumour burden for each treatment group measured by bioluminescence imaging (BLI) (n=6 mice per group). (**e**) Survival curves for all PDAC and TNBC tumour bearing mice treated with different CAR-T/T cell modifications. (**f**) and (**g**) ROR1 and HSD11B1 mRNA and protein expression at *in vivo* endpoint analyses from PDAC and TNBC tumours. (**h**) Association between ROR1 and HSD11B1 mRNA expression in PDAC tumours. (**i**) HSD11B1 mRNA expression after treatment with increasing effector-to-target ratios of CAR-T cells that secrete TNFα and IFNγ in PANC-1 cells. Statistical analyses were performed using multiple paired *t*-tests (**a,b**), mixed-effects analysis (**d**), log-rank (Mantel-Cox) test (**e**), one-way ANOVA with Dunnett’s test for multiple comparisons (**f,g,i**), and Pearson-correlation (**h**) and were considered significant if *p*-value was \**p*<0.05,\*\**p*<0.01,\*\*\**p*<0.001, \*\*\*\**p*<0.0001; ns, not significant; ELISA, enzyme-linked immunosorbent assay.

We obtained progressing PDAC and TNBC tumours at the experiment endpoint in all treatment groups and found that *ROR1* transcript and ROR1 protein expression were higher in mice that had received ^hGR-KO^ROR1 CAR-T cells, and to a lesser extent also non-edited ROR1 CAR-T cells, compared to UTD T cells **(Fig. 7f,g)**. This was surprising, as antigen downregulation or even antigen loss is frequently observed in tumours that persist after CAR-T cell therapy.^32,33^ Furthermore, there was a strong correlation between *HSD11B1* and *ROR1* expression in tumour lesions **(Fig. 7h and Extended Data Fig. 11i)**. We also noted that there was a strong correlation between *HSD11B1* expression in tumour lesions and DOC concentration in mouse serum **(Extended Data Fig. 10j)**, supporting that HSD11B1 was catalytically active and fueled the autocrine signaling pathway that augments ROR1 expression with recycled GCs in cancer cells in both xenograft models in vivo.

Of particular interest, we found that *HSD11B1* transcripts in tumour cells were higher after treatment with ^hGR-KO^ROR1 CAR-T cells and non-edited ROR1 CAR-T cells compared with UTD T cells **(Fig. 7f,g and Extended Data Fig. 11g,h)**. TNFα and other immune cell cytokines have been reported to induce HSD11B1 in cancer cells,^32^ suggesting that PDAC and TNBC may respond to immunological stress from CAR-T cells with this mechanism, resulting in augmented GC recycling and subsequent immune-suppression. Indeed, when we exposed PANC-1 cells and MDA-MB231 cells to increasing numbers of ^hGR-KO^ROR1 CAR-T cells and different doses of TNFα *in vitro*, there was a gradual increase in *HSD11B1* transcripts in tumour cells that we obtained during the anti-tumour response **(Fig. 7i and Extended Data Fig. 11k)**.

Taken together, these data show that ^hGR-KO^ROR1 CAR-T cells are resistant to the immunosuppression imparted by GC recycling and release in PDAC and TNBC, and confer potent anti-tumour reactivity in xenograft models *in vivo*. CAR-T cell cytokines that are released during the anti-tumour response, trigger a further increase in HSD11B1 activity in PDAC and TNBC cells that augments GC recycling and release, but also ROR1 expression due to autocrine GC signaling. Collectively, the data obtained in the setting of PDAC and TNBC provide examples of autocrine GC signaling that regulates ROR1 expression in steroidogenic non-endocrine cancers, and show that ^hGR-KO^ROR1 CAR-T cells are resistant to the paracrine effects of recycled GCs **(Extended Data Fig. 12 – Graphical Abstract)**.

## DISCUSSION

We assessed the autocrine and paracrine effect of endogenous GCs that are either produced *de novo* or through recycling on CAR-T cells directed against the oncogenic driver and cell surface antigen ROR1. Our data show that *de novo* GC production in ACC fuels an autocrine GC signaling loop in which hGR and STAT3 physically interact and are jointly required for ROR1 upregulation. We used inhibitors of GC production (e.g. mitotane) and hGR signaling (e.g. relacorilant), and CRISPR-Cas9 *hGR* knockout to demonstrate a direct mechanistic link between autocrine GC signaling and ROR1 transcription in ACC. Thus, the same tumour-derived hormonal signal exerts opposing effects across cellular compartments: autocrine GC signaling increases ROR1 antigen abundance in cancer cells, whereas paracrine signaling suppresses therapeutic T cells, mechanistically coupling therapeutic antigenicity to immune evasion. To resolve this conundrum, we engineered GC-resistant ROR1 CAR-T cells through CRISPR-Cas9 *hGR* knockout to block the paracrine effect of ACC-derived GCs on T cells. ^hGR-KO^ROR1 CAR-T cells maintained their phenotype and function in the setting of GC excess and vastly outperformed non-gene-edited ROR1 CAR-T cells in models of ACC *in vitro* and *in vivo*. Importantly, this principle extends beyond autonomous steroidogenesis in endocrine cancer: PDAC and TNBC recreate the same GC-hGR/STAT3-ROR1 axis through HSD11B1-mediated local GC recycling, where ^hGR-KO^ROR1 CAR-T cells similarly show superior antitumour activity.

Our findings further reveal an asymmetry between endogenous and engineered T cell immunity. GC-rich environments suppress T cell function and have been reported to impair peptide-MHC-dependent immune recognition,^34,35^ whereas CARs recognize intact surface antigens independently of MHC.^20^ This distinction may be particularly relevant in GC-rich tumours, where preserved or enhanced surface antigen expression can remain therapeutically accessible to CAR-T cells despite local immunosuppression.

Autocrine signaling in endocrine cancers and hormonally-active cancers that derive from non-endocrine tissues has thus far received limited attention in basic and translational cancer research. Prior studies have reported on autocrine signaling through vascular endothelial growth factor and heat shock proteins that promote proliferation and progression in gastric cancer.^36,37^ We show that GC-activated hGR physically interacts with STAT3 and that both factors are jointly required to sustain GC-dependent ROR1 expression and that this hGR/STAT3 complex can be disrupted by GC inhibitors.^28,38^ Earlier studies have reported that in TNBC, the hGR synergizes with STAT3 to instruct the transcriptional program that leads to a more aggressive basal-like tumour phenotype. STAT3 has been shown to be constitutively expressed in several types of cancer and has been positively correlated with immune cell infiltration.^28,38^

ROR1 is an oncofetal antigen involved in cell polarization and migration during embryonic development and tissue formation.^39,40^ In lymphoma and several solid cancers, ROR1 has been implicated in oncogenic signaling, and high ROR1 expression is frequently associated with cancer progression and development of metastases.^41–44^ A recent study has reported that administration of dexamethasone promotes the formation of metastases in preclinical models of breast cancer and reported increased expression of hGR and ROR1 after dexamethasone treatment.^45^ Another study reported that GCs induce differentiation and chemoresistance in ovarian cancer, and that dexamethasone increases ROR1 expression in ovarian cancer cell lines. The authors postulated an indirect mechanism between hGR signaling and ROR1 expression, based on the observation that ROR1 downregulation did not affect hGR expression.^46^ Our study reveals that GC-activated hGR signaling is directly linked to ROR1 transcription in ACC, as demonstrated by the antagonistic effect of GC inhibitors and the agonistic effect of GCs on ROR1 expression. We confirmed this antagonism using data obtained from our clinical cohort of patients with ACC^GC+^ treated with mitotane. Our clinical data also showed that high ROR1 expression in ACC correlates with advanced stage at diagnosis, aggressive disease course, and worse clinical outcome compared to ACC^GC−^. In previous work, WNT5A has been shown to be a ligand for ROR1 and to activate ROR1 signaling, which in turn augments ROR1 expression through STAT3.^47–49^ Upon treatment with GC inhibitors, we detected increased WNT5A in both ACC^GC+^ and ACC^GC-^ in our experiments.

Taken together, these data support our working model in which autocrine GC signaling induces ROR1 as an oncogenic driver that augments and perpetuates the aggressive phenotype of ACC^GC+^ (**Extended Data Fig. 11, Graphical Abstract**). The hGR/STAT3 complex therefore represents a node of dual therapeutic relevance: its disruption by GC inhibitors suppresses ROR1 and relieves oncogenic signaling, but simultaneously destroys the immunotherapeutic target antigen – a mechanistic tension that is resolved by restricting GR ablation to the CAR-T cell product rather than the tumour.

The majority of ACC lesions is lymphocyte-depleted, with negligible antitumour reactivity from endogenous adaptive and innate immune cells, an effect attributed to the immunosuppressive properties of ACC-derived GCs, which are produced *de novo* from cholesterol in the steroidogenic pathway.^14,50^ We reasoned that additional genetic engineering may be necessary to make CAR-T cells resistant to the paracrine, immunosuppressive effects of GCs.^6,51,52^ We demonstrate that ROR1 CAR-T cells, when shielded from GC excess in ACC^GC+^, TNBC and PDAC maintain their function and produce effective antitumour responses in mice. We chose immunodeficient NSG mice in all our xenograft models to allow attributed intra-tumoural GC production with confidence to tumour-cell-autonomous steroidogenesis or HSD11B1-mediated recycling because other endogenous immune and stromal populations that independently express GC-metabolizing enzymes are absent. Hence, the tumour model provides the mechanistic resolution necessary to establish a direct causal link between cell-autonomous GC production, ROR1 antigen amplification, and differential CAR-T cell efficacy. Future studies in humanized mouse models will be valuable for evaluating the interaction between ^hGR-KO^ROR1 CAR-T cells and endogenous immune effectors in a more physiologically complete setting, complementing the mechanistic clarity that the immunodeficient model provides.

Exogenous synthetic GCs such as dexamethasone are frequently administered to control cytokine release syndrome (CRS), immune-cell associated neurotoxicity syndrome (ICANS), or hematological toxicity in patients that have received CAR-T cells.^53,54^ Several studies reported a negative impact of exogenous GCs on the clinical outcome of CAR-T therapy.^6^ We anticipated that GC therapy would still be effective in preventing and treating these toxicities as GCs act on non-CAR-modified adaptive and innate immune cells and on endothelial cells that are involved in the respective adverse outcome pathways.^55,56,57,58^ The development of GC-resistant T cells has been explored in the context of treating viral infections and graft-versus-host disease in bone marrow transplant recipients.^59^ A prior study also developed IL13Rα2 CAR-T cells with hGR knockout using zinc-finger gene editing for glioblastoma therapy, where exogenous synthetic GCs are used to treat cerebral edema.^60^ However, the inclusion of safety switches or depletion markers such as EGFRt in this study into GC-resistant CAR-T cells is warranted to provide physicians with effective means to limit or terminate their activity in the event of adverse events.^61,62^

Endogenous GCs are increased in many cancer patients as part of the chronic stress response associated with the malignancy.^7^ Such endogenous GCs may originate from *de novo* production through steroidogenesis or from recycling of inactive metabolites like cortisone into active cortisol through the enzyme HSD11B1.^11,63^ The release of endogenous GCs has been reported in several non-endocrine cancers, including pancreatic,^64^ colorectal, gastric, and esophageal cancers.^11,45,65,66^ Transcriptomic analyses revealed that HSD11B1 expression is enriched in malignant epithelial cells, establishing tumour-cell-intrinsic GC recycling as the primary driver of intra-tumoural cortisol concentrations in this entity. Genomic copy-number gains affecting HSD11B1 were frequent in several tumour entities, suggesting that genomic alterations may further contribute to sustained HSD11B1 expression in selected cancers. However, this further underscores the clinical necessity of a GC-resistance approach that operates independently of HSD11B1 enzymatic activity.

We show that GC recycling and release in TNBC and PDAC, respectively, diminishes the efficacy of ROR1 CAR-T cells and that GC-resistant ROR1 CAR-T cells confer potent anti-tumour reactivity *in vitro* and *in vivo*. When we placed PDAC and TNBC under therapeutic pressure from ROR1 CAR-T cells, we observed a further increase in HSD11B1 activity and GC recycling, demonstrating a CAR-T-cell-induced stress response in both types of cancer. This adaptive induction of HSD11B1 by CAR-T cell-derived cytokines – most notably TNFα – represents a previously undescribed immune evasion mechanism whereby non-endocrine tumours actively amplify their GC recycling capacity in response to immunological stress. This creates a self-reinforcing resistance circuit: immune pressure increases GC recycling and suppression of non-edited CAR-T cells, whereas ^hGR-KO^CAR-T cells remain protected while simultaneously benefiting from GC-driven ROR1 amplification. Thus, the adaptive response used by the tumour to escape immune attack reinforces the therapeutic vulnerability exploited by GC-resistant CAR-T cells. Excessive GC release may foster the migration of suppressive immune cells and diminish the engagement of other adaptive and innate immune cells into the anti-tumour response.^3,9,10,16^

Taken together, our findings identify endogenous tumour steroid metabolism as an immune-regulatory circuit that couples immune suppression to therapeutic antigenicity. Selective hGR-KO in CAR-T cells uncouples these opposing effects by preserving GC-driven ROR1 amplification in cancer cells while protecting therapeutic T cells from GC-mediated suppression. This allows GC-resistant CAR-T cells to exploit a tumour-intrinsic endocrine immune-evasion mechanism as a therapeutic vulnerability. Although exemplified here by ROR1, identification of additional GC-regulated surface antigens may extend this principle across hormonally active cancers. More broadly, tumour-derived immunosuppressive signals may not necessarily need to be eliminated if their effects can instead be selectively uncoupled across cellular compartments and exploited by engineered immune cells.

## METHODS

### Patient samples

Tumour samples and nAG were obtained local biobank of the Division of Endocrinology & Diabetes at the University Hospital Wuerzburg, Germany, as part of the European Network for the Study of Adrenal Tumours (ENSAT) project. All patients provided prior written informed consent as part of the ENSAT ACC registry.^67^ The study followed the principles of the declaration of Helsinki, the good clinical practice guidelines and was approved by the ethics committee of the University of Wuerzburg (#88/11). In total, we included 197 tumour samples (n=149 primary tumours, n=14 local recurrences, n=34 ACC metastases) from 149 individual patients with histologically confirmed ACC and 22 nAGs were obtained from the biobank. The assessed clinical and histopathological characteristics included sex, age at diagnosis, tumour size, resection status, Ki67 proliferation index, Weiss score, ENSAT classification tumour staging, hormone secretion profile, steroid inhibitor treatment (metyrapone, ketoconazole, and mitotane) prior to surgery and the presence of local recurrences and distant metastases **(Table 1)**. Single cell transcriptomic data were analyzed from the RNA sequencing data set from Giordano et al.^21^ Overexpression of ROR1 as compared to nAGs was confirmed by qRT-PCR in a second, independent patient cohort consisting of 62 ACC samples from 51 individual patients (n=47 primary tumours and n=15 metastases) and 13 nAGs. ROR1 expression at protein level was finally confirmed by chromogenic immunohistochemistry in 135 ACC formalin-fixed paraffine-embedded (FFPE) tissue samples (n=102 primary tumours, n=14 local recurrences and n=19 metastases) and 9 nAGs. Endocrine workup confirmed autonomous glucocorticoid (GC) excess in 61.7% of all patients diagnosed by means of pathological 1 mg dexamethasone test in the presence of suppressed adrenocorticotropic hormone (ACTH).^67^ Completeness of surgical resection of the primary tumour was based on negative surgical, pathological and imaging reports (R0) for any residual malignant tissue. At the time of diagnosis and during follow up examinations at an interval of 3-6 months, the presence of local recurrences or metastases was evaluated by computed tomography of chest and abdomen.

### Cell lines, cell culture and 3D spheroid models

The ACC cells - NCI-H295R (from ATCC),^26^ JIL-2266 (in house),^27^ CU-ACC1 and CU-ACC2 (by Katja Kiseljak-Vassiliades)^25^ - were cultured as previously described. Non-adrenal cell lines including MDA-MB231 (TNBC) and PANC-1 (PDAC) were cultured in RPMI-medium (Gibco) supplemented with 9% FCS and 1% penicillin-streptomycin (pen-strep). Besides dexamethasone (Sigma-Aldrich) treatment (72 hours), cells were treated with different drugs dissolved in DMSO (metyrapone, ketoconazole, relacorilant & mitotane) for 48 hours. 3D cell culture was performed with 15% methylcellulose for 3 weeks.

### RNA extraction, silencing and RNA-nanostring nCounter analysis

RNA was extracted by using the Maxwell® RSC simply RNA Tissue Kit (Promega) according to the instructions of the manufacturer. RNA quality and quantity were determined using a Nanodrop 1000 (Thermo Fisher). hGR gene silencing was achieved using ACCELL siRNA. Digital multiplexed NanoString nCounter analysis system (NanoString Technologies)-based gene expression profiling was performed on 40 ng total RNA from each sample according to the manufacturer’s instructions. Nanostring RNA analysis of 785 immune cell and exhaustion-related human genes was performed using the nCounter® Immune Exhaustion Panel on the nCounter® Analysis System. Analysis and normalization of the raw Nanostring data were performed using nSolver Analysis Software v1.1 (Nanostring Technologies).

### Quantitative real-time-PCR (qRT-PCR)

Gene mRNA expression analysis was investigated by quantitative real-time polymerase chain reaction (qRT-PCR). RNA was isolated as described in the previous paragraph. Reverse transcription of mRNA into cDNA was performed by using the QuantiTect Reverse Transcription Kit (Qiagen) according to manufactureŕs recommendations. mRNA expression was analyzed in duplicates and normalized to α-Tubulin. Gene amplification during qRT-PCR was performed on a CFX96 real-time thermocycler (Bio-Rad) and Bio-Rad CFX Manager 2.0 software in a 50 µl reaction consisting of 5.0 ng cDNA, 300 nM gene-specific forward and reverse primers and 25 µl Power SYBR Green PCR Master Mix (Applied Biosystems). Cycling conditions were 95°C for 10 minutes (min) followed by 40 cycles of 95°C for 15 s, 60°C for 1 min. The cycle threshold (Ct) was determined using SDS software v2.2.2 (Applied Biosystems) and the level of gene expression calculated by using the ΔCt method (2−(ΔCt)).

### RNAscope single cell analysis

RNAScope is a custom RNA in situ hybridization solution (Advanced Cell Diagnostics) that allows the staining of RNA fragments on FFPE tissue slides. Staining has been performed according to manufactureŕs protocol and extensively elucidated priorly.^68^ Three pictures of representative areas of each slide were taken with the Leica Aperio Versa brightfield scan-ning microscope (Leica) at 40x magnification. For scoring the slides, optional image analysis algorithm ‘RNA ISH v1’ of the Aperio ImageScope software v.12.x (Leica) has been used on the entirety of the pictures. This algorithm automatically detects and counts the number of RNA molecules (each brown stained spot is one molecule of RNA) and the number of cells (by detecting the hematoxylin-stained nuclei in a defined area. Thresholds for the detection were manually adjusted for a high-fidelity assessment of the signal. We used the ratio of RNA molecules per cells for each slide to quantify ROR1 gene expression (Hs-ROR1, #402831, Advanced Cell Diagnostics).

### Chromogenic and immunofluorescence immunohistochemistry (IHC) and H-scores

Chromogenic and fluorescence immunohistochemical staining on human FFPE tumour slides has been performed as already reported.^14,68^ The primary antibodies that have been used were the following: ROR1 (4A5, BD Pharmingen), CYP11B1 and CYP11B2 (provided by Celso Gomez-Sanchez),^69^ SF-1 (N1665, R&D Systems), Ki67 (MIB-1, Dako), CD31 (orb10314, Biobyrt), CD3 (ab699, abcam), p53 (DO-7, Bio-Rad), hGR (D4X9S, Cell Signaling), Stat3 (D1B2J, Cell Signaling), pStat3 (D3A7, Cell Signaling), Inhibin-α (R1, Bio-Rad). Microscopic assessment and H-score evaluation was performed using Aperio Leica scanner and Aperio Leica ImageScope software analysis (Leica).

### Direct stochastic optical reconstruction microscopy (*d*STORM)

Purified anti-human ROR1 antibody was purchased from Biolegend (Cat. 357802). Buffer exchange was carried out using Zeba Spin Desalting Columns and antibodies were conjugated with Alexa Fluor 647 using the AF647-NHS ester kit (Thermo Scientific) according to the manufacturer’s protocols. The concentration of the final conjugated antibody solution was determined on a Tecan Spark. 8-well chambered cover glass chambers (Cellvis) were coated with 0.02 mg/ml PLL or PDL. Tumour cells were harvested with a cell scrapper, resuspended in FACS buffer. Fc-Receptor blockade was carried out using 1:9 diluted TruStain FcX, followed by ROR1 staining using 10 µg/ml ROR1-AF647 antibody for 30 min at 4°C in the dark. Background signal from unspecific antibody binding was removed by sequen-tial wash steps. Cells were plated in coated cover glass chambers and allowed to adhere for 15 min on ice and fixated 3% formaldehyde and 0.25% glutaraldehyde in PBS for 15 min. Fixation solution was removed and washed 3x with PBS. For direct stochastic optical reconstruction microscopy (dSTORM), an ONI Nanoimager S was used. For *d*STORM measurements, imaging buffer (100 mM cysteamine hydrochloride in PBS, pH 7.4) was added to cover slides. Images were taken in total inner reflection fluorescence (TIRF) mode. Pixel size was 117 nm. Typically, 15000 frames at 10 ms exposure time were taken per movie with a laser power at 640 nm wavelength of ∼3.5 kW/cm². *d*STORM image reconstruction was performed with rapidSTORM 3.3.^70^ Drift correction was performed using the linear drift correction tool of rapidSTORM 3.3. Selection of cell region of interests (ROIs) was performed with Napari and cross checked with widefield images. For cluster analysis, a custom localization analysis tool LOCAN was used, which employs a density-based spatial clustering of applications with noise (DBSCAN) algorithm.^71^ Cluster parameters of DBSCAN were minPoints = 3 and epsilon = 20 nm.

### Western Blot analysis

For Western Blot (WB) analysis, proteins were extracted from fresh-frozen ACC tissues and cells as previously described.^72^ Protein concentration was quantified as previously described.^73^ The primary antibodies that have been used were the following: ROR1 (D6T8C, Cell Signaling, #16540S), Wnt5A (Cell Signaling, #2392S), hGR (D4X9S, Cell Signaling, #47411S), Stat3 (D1B2J, Cell Signaling, #30835S), pStat3 (D3A7, Cell Signaling, #9145S).

### Duolink® Proximity Ligation Assay (PLA®)

ACC tumour samples and 3D cell culture spheroids have been formalin-fixed, paraffine-embedded and applied on a microscopic slide. Duolink® Proximity Ligation Assay (PLA®) has been performed according to the instructions of the manufacturer. In brief, tissue slides have been initially blocked for 1 hour at 37°C in a heated chamber with the Duolink® blocking solution (Merck). After multiple washing steps, cells were incubated successively with PLA probes, ligation solution and amplification solution at 37°C (Duolink© In Situ PLA® Probe Anti-Mouse MINUS & Anti-Rabbit PLUS). Antibodies have been the same from IHC. Samples have been covered with DAPI, cover slips and sealed with transparent nail polisher. Images were examined using Aperio Leica scanner (Leica).

### (Quantitative) flow cytometry (FACS) analysis and cell phenotyping

Cells were washed with FACS buffer before staining. Staining was performed in FACS buffer for 20 min at 4°C. Cells were then washed once with FACS buffer and analyzed on a BD FACS Canto II. FACSDiva software (BD) was used for data collection and FlowJo software (BD) was used for data analysis. T cells were detected for CAR expression using an tEGFR antibody. The following antibodies were used for cell staining: ROR1 (AF647, clone 2A2, BioLegend), CD45 (VioBlue, clone REA747, Miltenyi), CD4 (PeVio770, clone REA, Miltenyi), CD8 (APC-Cy7, clone SK1, BioLegend), tEGFR (APC, clone AY13, Bio-Legend), PD-1 (PE, clone PD1.3.1., Miltenyi), Lag3 (PerCP-Cy5.5, clone 11C3C65, Bio-Legend), CTLA-4 (PeCy7, clone L3D10, BioLegend), Tim-3 (CD366), APC-Cy7, clone F382E2, BioLegend), CD45RO (FITC, clone UCHL1, BioLegend), CD45RA (PE, clone T6D11, Miltenyi), CD62L (PerCP-Cy5.5, clone DREG-56, BioLegend), CD25, PeCy7, clone BC96, BioLegend), CD69 (APC-Cy7, clone FN50, BioLegend). Quantitative flow cytometry was performed using PE-labeled antibodies and the BD-Quantibrite PE-Bead Assay (BD Biosciences).

### Liquid chromatography-tandem mass spectrometry (LC-MS/MS) analysis

Steroid hormones in cell culture supernatants of ACC cell lines and mouse blood were quantified using a liquid chromatography-tandem mass spectrometry system (QTRAP 6500+, SCIEX®) including an Agilent 1290 UHPLC (G4226A autosampler, infinityBinPump, G1316C column-oven, G1330B thermostat). The MassChrom-Steroids in Serum/Plasma® IVDR conform kit (Chromsystems®) was used for steroid measurements. 15 steroid hormones in the positive MRM-Mode (aldosterone and DHEAS in the negative mode) were quantified via corresponding isotope-labeled standards according to the manufacturer’s instruction. 500 μl serum/plasma/supernatants were utilized for offline solid phase extraction (SPE) and finally 15 μl were used for LC-MS/MS. Raw data were processed by Analyst® Software (1.6.3) via 6 point calibration curve and 1/x weighting. Periodic participation in ring trails and commercial quality controls ensured the correctness of measurements for the steroids.

### T cell isolation, activation, expansion and culture

T cells were isolated from PBMCs by negative selection using CD4^+^ and CD8^+^ T cell isola-tion kits (Miltenyi), and have been expanded and activated using anti-CD3/anti-CD28 bead stimulation (Invitrogen) prior to nucleofection. They were cultured in T cell medium (RPMI-1640 supplemented with 9% human serum, 1% pen-strep, 0.1% GlutaMax) and 50 U/mL recombinant human interleukin (IL)-2 (Proleukine). Patient-derived and healthy donor CD4^+^ and CD8^+^ T cells were isolated from peripheral blood by negative selection. CAR-transduced T cells were enriched using biotinylated anti-EGFR mAb and anti-biotin beads (Miltenyi), prior to expansion using a rapid expansion protocol.^74,75^

### Pan-cancer tumour-normal meta-cell atlas

Normalized scRNA-Seq date were download from Kang et al. covering 30 different cancer types across 104 datasets.^76^ A total of 4.9 million cells from 1,070 tumours and 493 normal samples from 999 donors were collected by the authores. To reduce the size of the single-cell data, we have generated meta-cells for each sample and cell type. A meta-cell represents the transcriptomes of a group of highly similar cells. We used the R package SuperCell v1.0 with additional parameter (γ = 15) to calculate approximately 300,000 meta-cells.^77^ Integration analysis was performed using Seurat v5.1.0^78^ for the **Figure 6b**. For each sample, the 2,000 most variable genes were estimated using the “vst” method of the FindVariableFeatures function. Mitochondrial, Immunoglobulin variable chain, T cell receptor and sex-specific genes were not included in the estimation of variable genes. We then used the SelectIntegrationFeatures function to select 2,000 genes that were repeatedly shown to be highly variable in multiple samples. Thereafter, the data were standardized with ScaleData, and principal component analysis (PCA) was performed on the standardized expression data of the variable genes using the function RunPCA. We used 25 principal components, which explained approx. 90% of the variance, to integrate all samples using the Harmony method (v1.2)^79^ with the Seurat wrapper function RunHarmony with additional parameters (dim.use = 1:25, group.by.vars = c(”study”)).

### CAR-T cell generation and CRISPR/Cas9-mediated glucocorticoid receptor (hGR) knockout

Two days after isolation and stimulation, 1.2×10^6^ CD4^+^ and CD8^+^ T cells were used for nucleofection adding 1µg/reaction of the sleeping beauty vector containing the R12.BBz CAR construct and 0.6 µg minicircle DNA encoding the SB100X transposase (Plasmid factory) to the transfection medium. HGR molecule expression was disrupted by CRISPR/Cas9-mediated gene editing. Single guide RNA for hGR was purchased from Integrated DNA Technologies and Cas9 protein from PNA Bio. Electroporation was performed according to the manufacturer’s protocol using the 4D-NucleofectorTM (Lonza, program EO-115). The electroporated cells (100 µl) were then transferred to a 48-well plate with 0.9 ml pre-warmed CTL medium and a half medium change with CTL supplemented with 100 U/ml recombinant human IL-2 was performed after 4h. CD3/CD28 Dynabeads were removed magnetically 4 days after transfection. Transfection efficiency was assessed the day after bead removal by flow cytometry via staining of the tEGFR transfection marker with biotinylated anti-tEGFR mAb (in-house).

### CAR-T cell proliferation assay

T cells were labeled with 0.2 mmol/L carboxyfluorescein succinimidyl ester (cell trace CFSE; life technologies), washed, and plated in triplicates with inactivated tumour cells at E:T ratio of 1:1 in cell culture medium without exogenous cytokines. After a 72 hours incubation, cells were labeled with CD8, CD4, tEGFR antibody. 7AAD (BD Biosciences) staining was conducted for live/dead cell discrimination. Samples were acquired by flow cytometry (FACS) to assess cell division by measuring CFSE dilution in living CAR^+^ T cells.

### CAR-T cell cytotoxicity and cytokine analysis

CAR-T cells were co-cultured with 50,000 tumour cells (2D cell culture) at the specified E:T ratios in RPMI medium on 96-well flat-bottom plates. For 3D cell culture, an E:T ratio of 0.25:1 was applied using 200,000 tumour cells per spheroid and well. Co-cultures were incubated at 37°C and analyzed after 24 hours. The basic analytic feature was used to quantify ATP to evaluate cell viability using the CellTiter Glo Assay (Promega) according to the manufacturer’s protocol. Specific lysis was calculated by subtracting T cell viability from the total viability of untreated tumours and normalizing it to UTD T cell reference. After 24 hours of co-culture, supernatants were collected and cytokine analysis was quantified by enzyme-linked immunosorbent assay (ELISA). Interferon (IFN)-γ and Interleukin (IL)-2 ELISA Kits were obtained from BioLegend. The analyses were performed according to the instructions of the manufacturer and absorbance was measured using a NanoQuant Infinite M200 Pro plate reader (TECAN).

### *In vivo* experiments and tumour engraftment in ACC xenograft mouse models

All animal studies were carried out according to the ARRIVE guidelines and animal experiment license that has been granted by the competent authorities and to the guidelines of the institutional animal care and use committee (University of Wuerzburg). Immunodeficient NOD.Cg-PrkdcSCID Il2rgtm1Wjl/SzJ (NSG) mice were purchased from Charles River. 8-week-old female mice were inoculated subcutaneously with 1×10^7^ NCI-H295R cells 14 days before T cell injection in 200 µl sterile PBS with 1% sterile FCS. Mice were blindly randomized and 1×10^6^ T cells were injected intravenously one time, 5 days after activation. Mice were monitored two to three times a week and tumour burden was quantified weekly using digital caliper measurement. Tumour area was calculated by multiplication of 3D tumour measurements using hemi ellipsoid formula (2/3 × π × length × width × depth). Mice were humanely euthanized when tumour burden reached the dropout criteria or showed signs of morbidity that were not in line with animal welfare or overreached a specific body condition score.

### Assessment of T cell expansion, persistence and exhaustion *in vivo*

To assess T cell expansion *in vivo*, blood samples were taken at day 10, 14, 21, 28 and when mice had to be removed from the experiment. Blood, tumour, spleen and bone marrow were collected, counted and analyzed using FACS analysis. All samples have been stained for human CD45, CD4, CD8, and CAR (tEGFR). Moreover, to assess T cell persistence, activation and exhaustion *in vivo*, at endpoint analyses intratumoural T cells were isolated by a tumour dissociation kit (Miltenyi) and stained for human CD45, PD-1, LAG3, CTLA-4, Tim-3 (CD366) and CAR (tEGFR). T cell subsets and memory have been evaluated by staining for human CD45, CD45RO, CD45RA, CD62L, CD25, CD69 and CAR (tEGFR).

### Statistical analysis

Statistical analyses were performed with Excel v.16.64 (Microsoft) and GraphPad Prism v.9.3.1 (GraphPad). Graphical schemes were designed with *Biorender.com*. Licenses are available on demand. Unless otherwise noted, analyses testing for significant differences between groups were conducted with unpaired two-tailed t-tests (when comparing two un-matched groups), paired t-tests (when comparing two matched groups), a one-way ANOVA with Dunnett’s test for multiple comparisons (when comparing more than two groups). We also used linear regression analysis to assess correlations between expression data and distinct clinical parameters. Survival curves were compared with the log-rank Mantel-Cox test. *In vivo* tumour growth was compared with a repeated measures ANOVA with correction for multiple comparisons. *In vivo* T cell proliferation and persistence were compared using the two-tailed Mann-Whitney test. P-values of less than 0.05 were considered statistically significant.

## Resource availability

### Lead contact, data and materials availability

This manuscript does not contain personal and/or medical information about an identifiable living individual. All data needed to evaluate the conclusions of this paper are either present in this paper and/or the supplementary materials. For sharing of the materials that have been used in this study, a material transfer agreement will be required. Requests for materials should be sent to M.F. & M.H.

## Acknowledgments & funding

The authors wish to acknowledge and thank Tanja Maier for supportive and constructive discussions on this project and manuscript, Alexandra Cirnu for providing microscope infrastructure, and Professor Celso Gomez-Sanchez for providing the anti-CYP11B1 antibody. The authors wish to thank Corcept Therapeutics for providing material (relacorilant) for research purposes and Hazel Hunt and Andreas Moraitis (Corcept Therapeutics) for advice. This study was supported through continuous sample collection and preparation by Sabine Kendl, Daniel Oppelt, Martina Zink and Felix Meindl, as well as collection and documentation of clinical data by Michaela Haaf at the University Hospital Würzburg. The authors wish to acknowledge the technical support of Quentin Deveuve, Richard Taylor and Linda Karschti in the *in vivo* studies. This project has received funding from the German Research Foundation (Deutsche Forschungsgemeinschaft (DFG), grants 314061271, INST 93/1033-1 FUGG – CRC/TRR 205 to MF and MK and FA-466-4-2 to MF, grant 338/1 2021-452881907 – CRC/TRR 338, project A02 to MH, and project Z02 to HE, project A05 to TN, project B05 to SD, grant 221/2 2021-X – CRC/TRR221, project A03 to MH and HE; Bavarian Cancer Research Center (Bayerisches Zentrum für Krebsforschung, BZKF), Leuchtturm Zelluläre Immuntherapien to MH, HE; German Cancer Aid (Stiftung Deutsche Krebshilfe), grant 70114707 (TransOnc Avant-CAR.de to MH) & grant 70115200 (Pre-clinical Drug Development Platform CAR FACTORY to MH); Federal Ministery for Education and Research (Bundesministrium für Bildung und Forschung, BMBF), grant 13N15987 (IMAGINE to MH, TN, PS), grant 01EN2306A (ROR2 CAR-T to MH), & grant 03ZU1111BA (SaxoCell to MH); Innovative Medicine Initiative 2 Joint Undertaking (JU), grant 853988 (imSAVAR) and grant 945393 (T2EVOLVE). The JU receives support from the European Union’s Horizon 2020 research and innovation program and EFPIA and JDRF International. Additional funding was provided by the patient advocacy group “Hilfe im Kampf gegen den Krebs e.V.” and the foundation “Forschung hilft – Stiftung zur Förderung der Krebsforschung an der Universität Würzburg” (to MF, JW, HE, MH).

## Author contributions

**M. P. Schauer:** Conceptualization, data curation, formal analysis, validation, investigation, visualization, methodology, project administration, writing–review and editing. **J. Weber:** Conceptualization, data curation, formal analysis, validation, writing–original draft, writing–review and editing. P. Spieler: Investigation, data curation, formal analysis, writing–review and editing. **B. Altieri:** provided clinical data, writing–review and editing. **F. Freitag:** Investigational support, writing–review and editing. **L. Gehrke:** Investigational support, writing–review and editing. **S. Sbiera:** Methodology, writing–review and editing. **S. Kircher:** Investigational support, pathology analyses, writing–review and editing. **M. Kurlbaum:** Investigation, Methodology, writing–review and editing. **M. Kroiss:** Funding acquisition, writing–review and editing. **K. Kiseljak-Vassiliades:** Provided cell lines, writing–review and editing. **M. E. Wierman:** Provided cell lines, writing–review and editing. **T. Nerreter:** Methodology, funding acquisition, writing–review and editing. **S. Danhof:** Funding acquisition, writing–review and editing. **H. Einsele:** Funding acquisition, writing–review and editing. **L-S. Landwehr:** Conceptualization, resources, supervision, funding acquisition, writing–original draft, project administration, writing–review and editing. **M. Fassnacht:** Conceptualization, resources, supervision, funding acquisition, writing–original draft, project administration, writing–review and editing. **M. Hudecek**: Conceptualization, resources, supervision, funding acquisition, writing–original draft, project administration, writing–review and editing.

## SUPPLEMENTARY INFORMATION

### TABLES

**Table 1.**
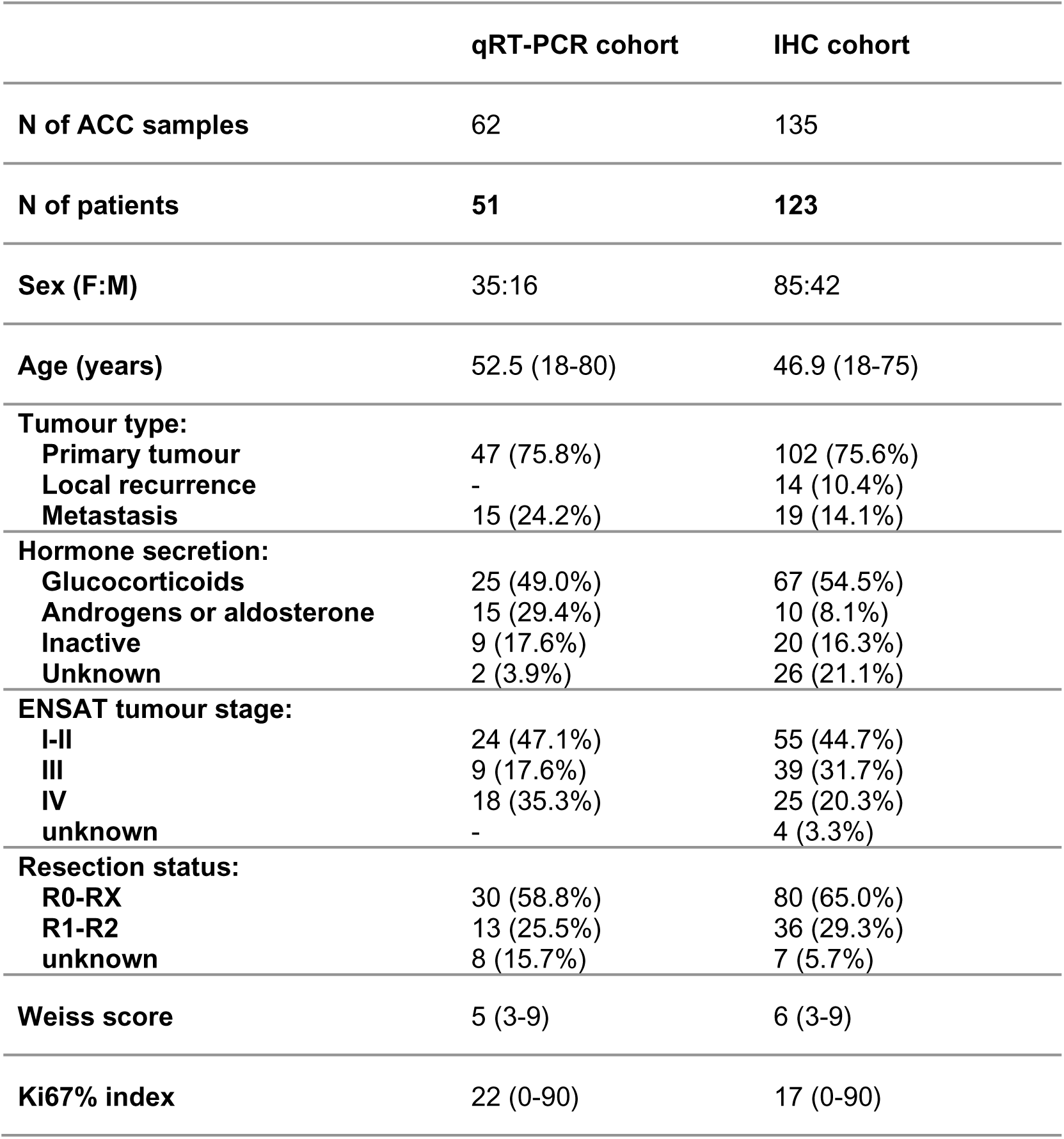
Adrenocortical carcinoma (ACC): patients’ and tumour characteristics. Data represent median values with ranges or total numbers. Tumour stage at the time of diagnosis according to the European Network for the Study of Adrenal Tumours (ENSAT) classification. In patients who experienced local recurrences or distant metastases during mitotane treatment, endocrine activity was classified according to the information available at primary diagnosis.

### Supplementary figures

**Extended Data Fig. 1.**
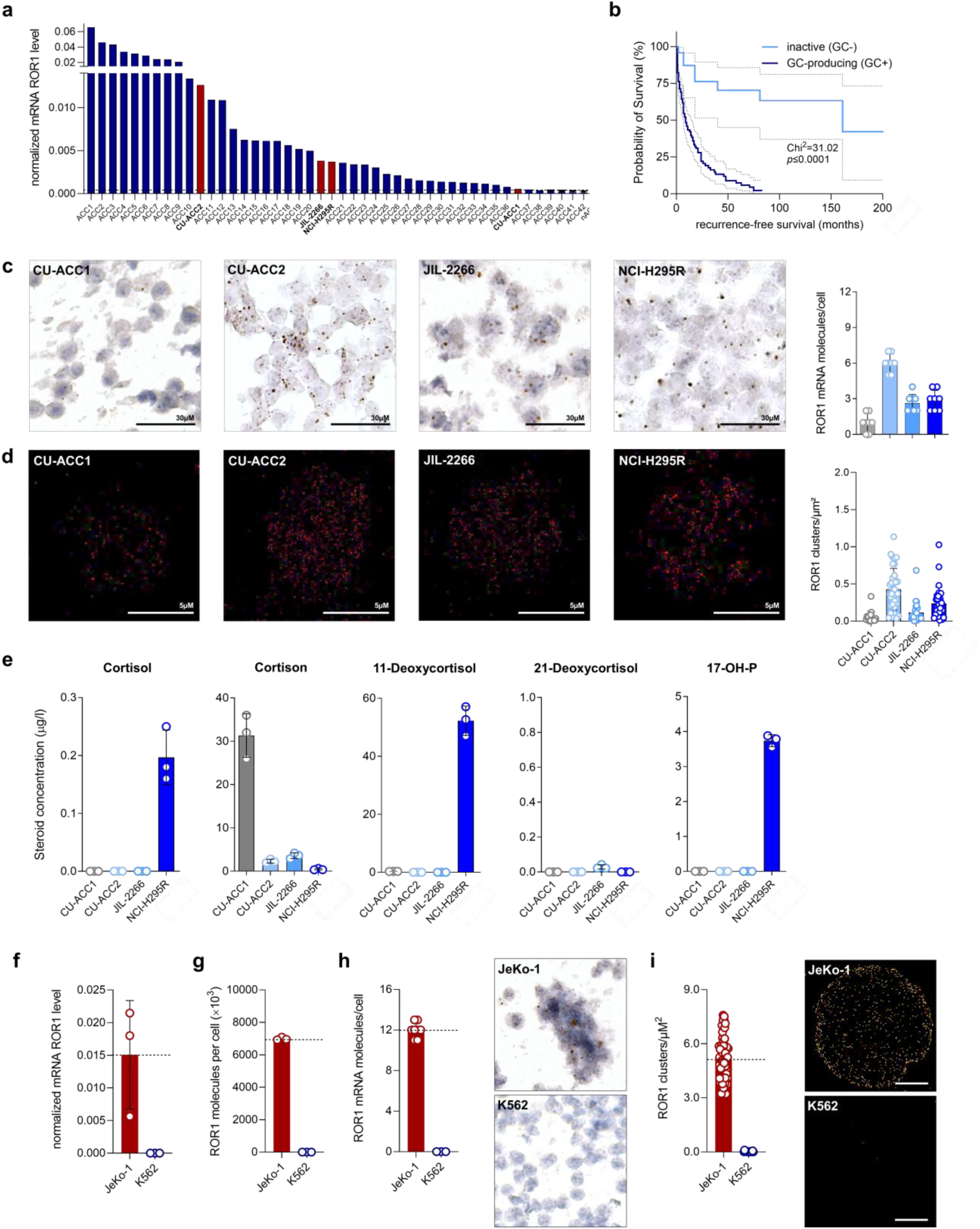
ROR1 expression in ACC cells and cell lines. (**a**) Waterfall plot comparing ROR1 mRNA expression in human ACC cell lines with ACC patients’ samples. (**b**) Survival curve comparing recurrence-free survival of ACC patients with and without glucocorticoid excess. (**c**) ROR1 mRNA molecules per cell of all four ACC cell lines assessed by RNAscope single cell analysis. Representative pictures are shown alongside. (**d**) ROR1 clusters/µm^2^ per basal plasma cell membrane of all four ACC cell lines assessed by single molecule super high-resolution microscopy (*d*STORM) analysis (n=30). Representative pictures are shown alongside. (**e**) Glucocorticoid secretion patterns of all four ACC cell lines assessed by LC-MS/MS analysis. (**f**) ROR1 mRNA expression of of JeKo-1 and K562 control cell lines assessed by quantitative real-time PCR (qRT-PCR). (**g**) ROR1 molecules per cell (×10^3^) of all four ACC cell lines assessed by quantitative flow cytometry (n=3). (**h**) ROR1 mRNA molecules per cell of JeKo-1 and K562 control cell lines assessed by RNAscope single cell analysis. Representative pictures are shown alongside. (**i**) ROR1 clusters/µm^2^ per basal plasma cell membrane of JeKo-1 and K562 control cell lines assessed by single molecule super high-resolution microscopy (*d*STORM) analysis (n=30).

**Extended Data Fig. 2.**
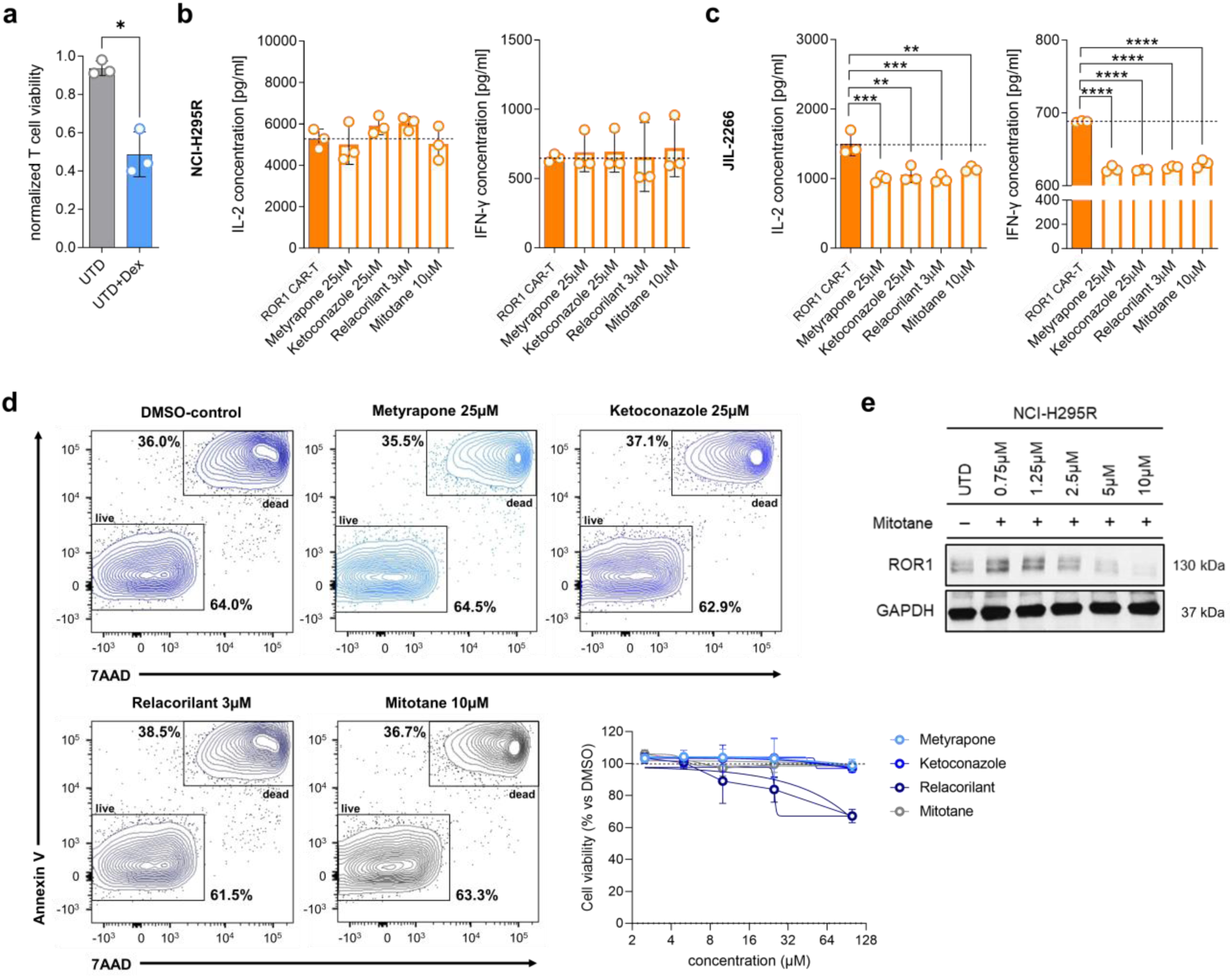
Diminished efficacy of ROR1 CAR-T cells against GC-inhibitor-treated ACC cell lines. (**a**) Cell viability of T cells after 72 hours of dexamethasone treatment. Cytokine secretion from ROR1 CAR-T cells in 3D cell culture models of (**b**) NCI-H295R and (**c**) JIL-2266 ACC cells after 48 hours of pretreatment with glucocorticoid inhibitors assessed by ELISA measurement. (**d**) Viability of CAR-T cells after 48 hours with different doses of the applied combination treatments (normalized to DMSO, n=3). Representative dot plots are shown alongside. (**e**) ROR1 protein expression after different doses of mitotane treatment assessed by western blotting. Statistical analyses were performed using one-way ANOVA with Dunnett’s test for multiple comparisons (**b,c**) and were considered significant if p-value was *p<0.05,**p<0.01,***p<0.001, ****p<0.0001; ns, not significant; hGR, human glucocorticoid receptor.

**Extended Data Fig. 3.**
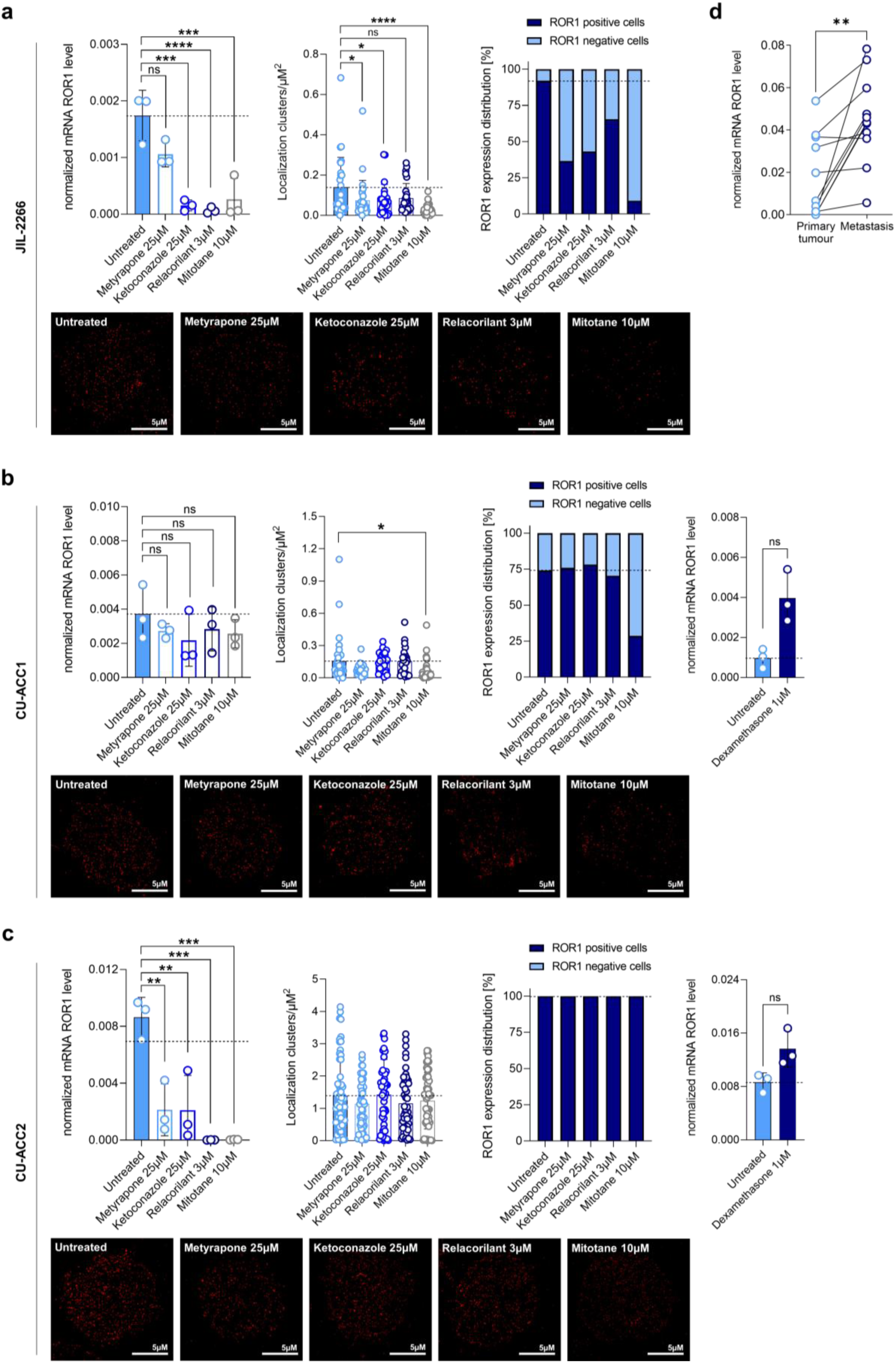
ROR1 downregulation after GC-inhibitor treatment in ACC cell lines. ROR1 mRNA and protein expression after glucocorticoid (GC) inhibitor treatment using metyrapone, ketoconazole, relacorilant and mitotane in (**a**) JIL-2266, (**b**) CU-ACC1 and (**c**) CU-ACC2 cell line assessed by qRT-PCR and *d*STORM analysis. (**d**) ROR1 mRNA and protein expression comparing untreated matched primary tumour samples with metastatic lesions of the same patient. Statistical analyses were performed using one-way ANOVA with Dunnett’s test for multiple comparisons (**a,b,c**) and paired *t*-test (**a-d**) and were considered significant if p-value was *p<0.05,**p<0.01,***p<0.001, ****p<0.0001; ns, not significant.

**Extended Data Fig. 4.**
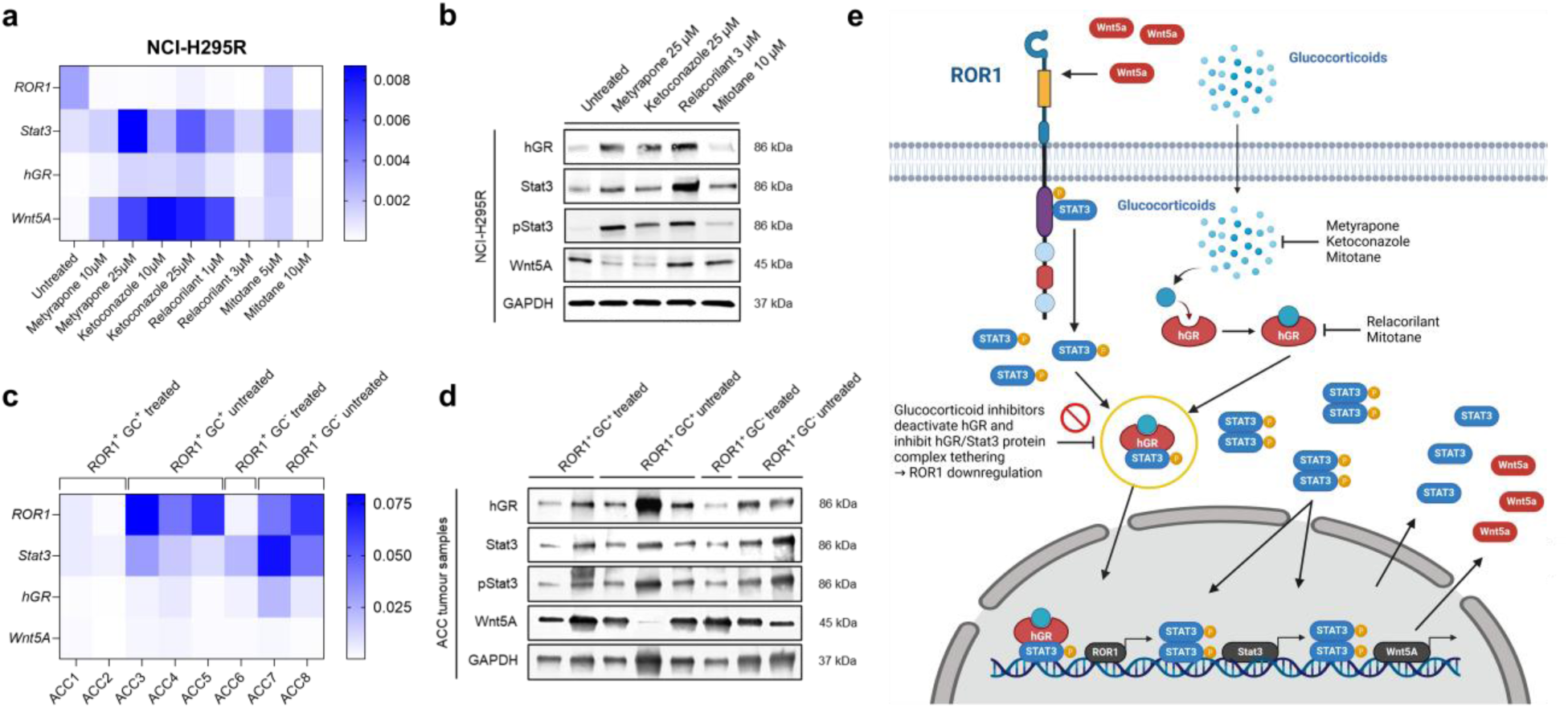
ROR1 regulating proteins act in a self sustaining loop upon GC inhibition. (**a**) RNA gene expression of key genes between the Wnt/ROR and the hGR/Stat3 pathway (n=3) in NCI-H295R cells after 48 hours of co-culture with pharmaceutical inhibitors of GC effector functions. (**b**) Western Blot analysis of Wnt5A, Stat3 and the hGR in NCI-H295R cells. (**c**) Expression of key genes between the Wnt/ROR and the hGR/Stat3 pathway (n=3) in primary ROR1 expressing ACC patient cells (ROR1^+^) that have been shown GC excess (GC^+^) or not (GC^−^) and either treated with GC inhibitors (treated) or left untreated (untreated) prior to surgery. (**d**) Western Blot analysis of Wnt5A, Stat3 and the hGR in primary ROR1 expressing ACC patient cells (ROR1^+^) that have been shown GC excess (GC^+^) or have been hormonally inactive (GC^−^) and either treated with GC inhibitors (treated) or left untreated (untreated) prior to surgery. (**e**) Scheme of the working model molecular pathway including the Wnt/ROR pathway and the investigated hGR/Stat3 interplay.

**Extended Data Fig. 5.**
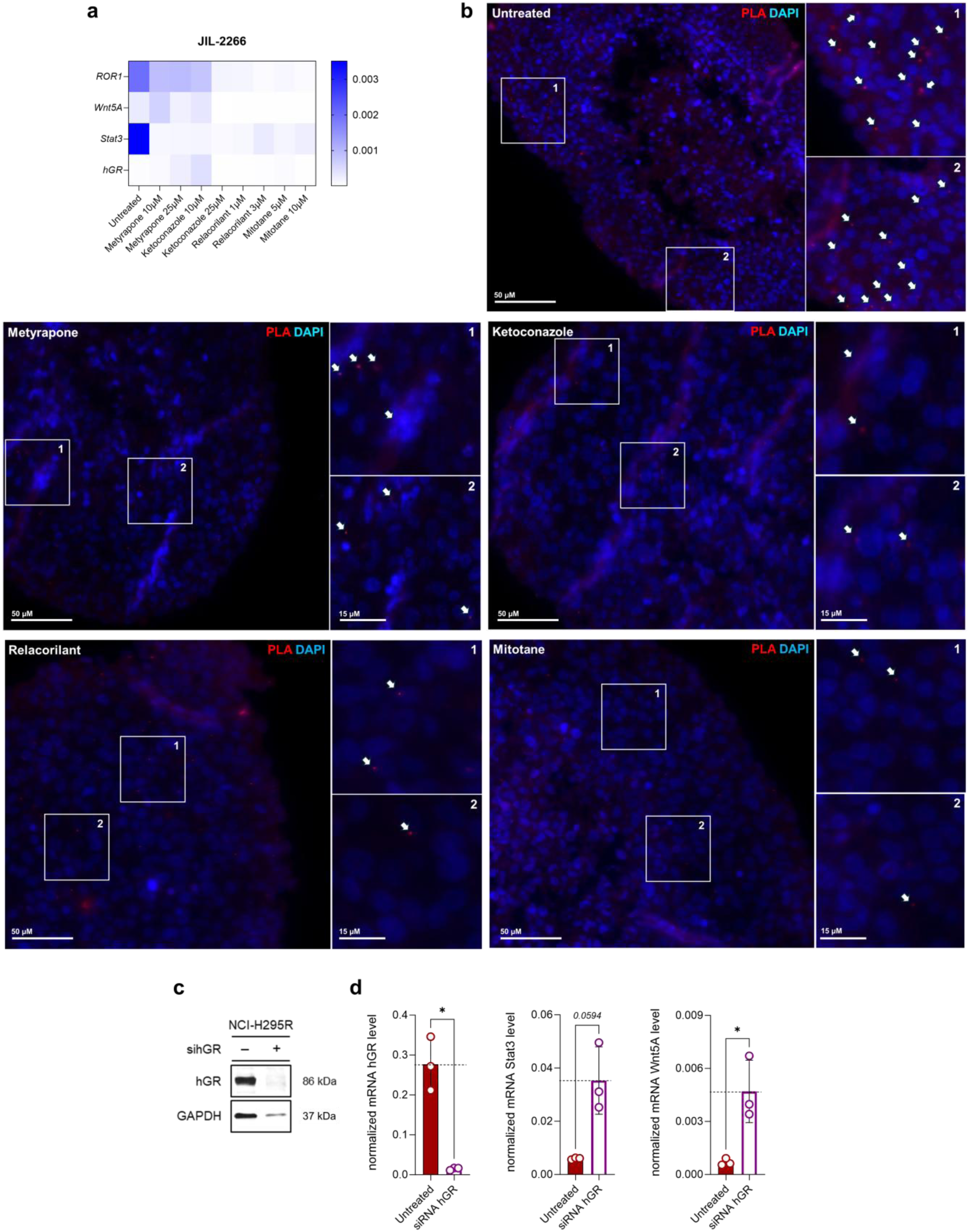
hGR gene silencing in ACC cell lines. (**a**) ROR1, Wnt5A, Stat3 and hGR mRNA expression before and after treatment with glucocorticoid (GC) inhibitors in JIL-2266 cell line. (**b**) Representative pictures of Duolink proximity ligation analysis (PLA) assessing hGR/Stat3 protein complex (PLA in red and DAPI in blue) in positive NCI-H295R cells in 3D cell culture before (untreated) and after 48 hours of treatment with different glucocorticoid (GC) inhibitors. (**c**) hGR protein expression after hGR gene silencing in NCI-H295R cells. (**d**) hGR, Stat3 and Wnt5A mRNA levels after hGR gene silencing. Statistical analyses were performed using paired *t*-tests (**a-c**) and were considered significant if p-value was *p<0.05,**p<0.01,***p<0.001, ****p<0.0001; ns, not significant; hGR, human glucocorticoid receptor.

**Extended Data Fig. 6.**
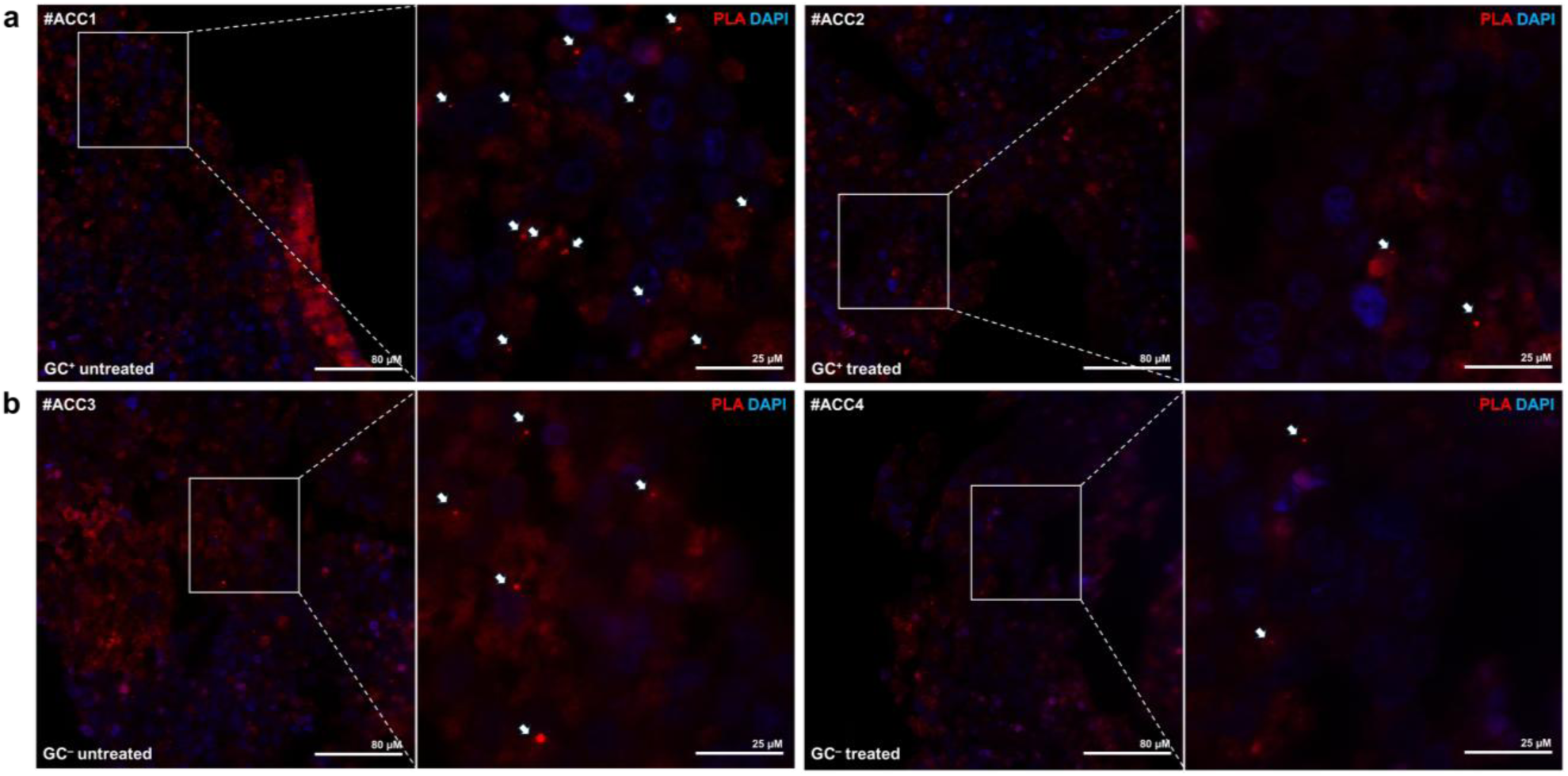
hGR/STAT3 complexes in primary ACC patient tumour samples. Representative pictures of Duolink proximity ligation analysis (PLA) assessing hGR/Stat3 protein complex (PLA in red and DAPI in blue) in primary ACC patient samples (a) with and (b) without GC excess before and after GC inhibitor treatment.

**Extended Data Fig. 7.**
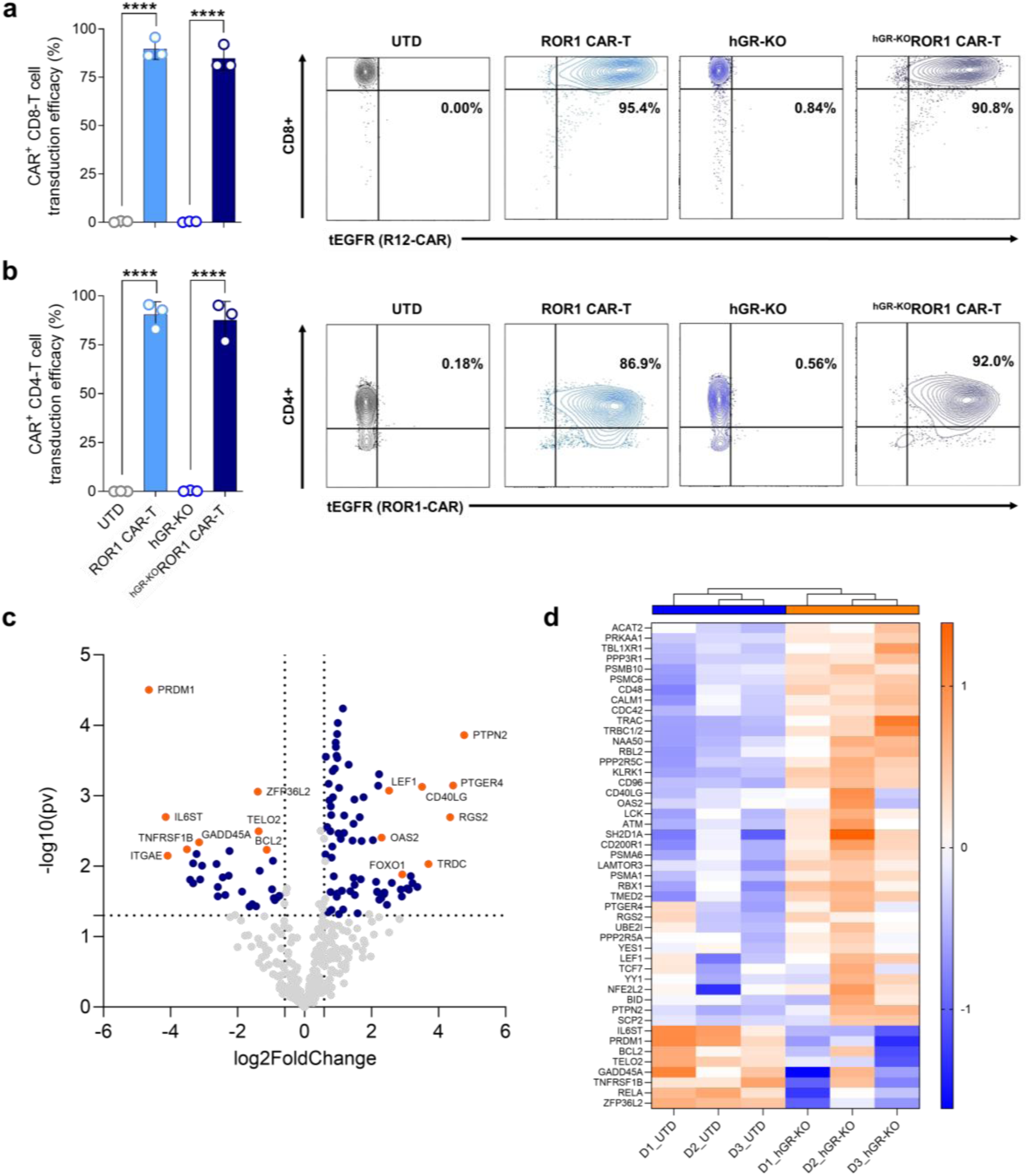
Phenotype and gene expression in GC-resistant ROR1 CAR-T cells. (**a-b**) CAR positive CD8 and CD4 T cells assessed by FACS analysis. Representative dot plots of ROR1 (R12.BBz) CAR expressing T cells and CD4+ and CD8+ T cell transduction efficacy are shown alongside. (**c**) Volcano plot and (**d**) corresponding heatmap of RNA Nanostring gene expression analyses comparing non-edited ROR1 CAR-T cells with ^hGR-KO^ROR1 CAR-T cells. Top 8 genes are annotated, horizontal line represents p-value <0.05.

**Extended Data Fig. 8.**
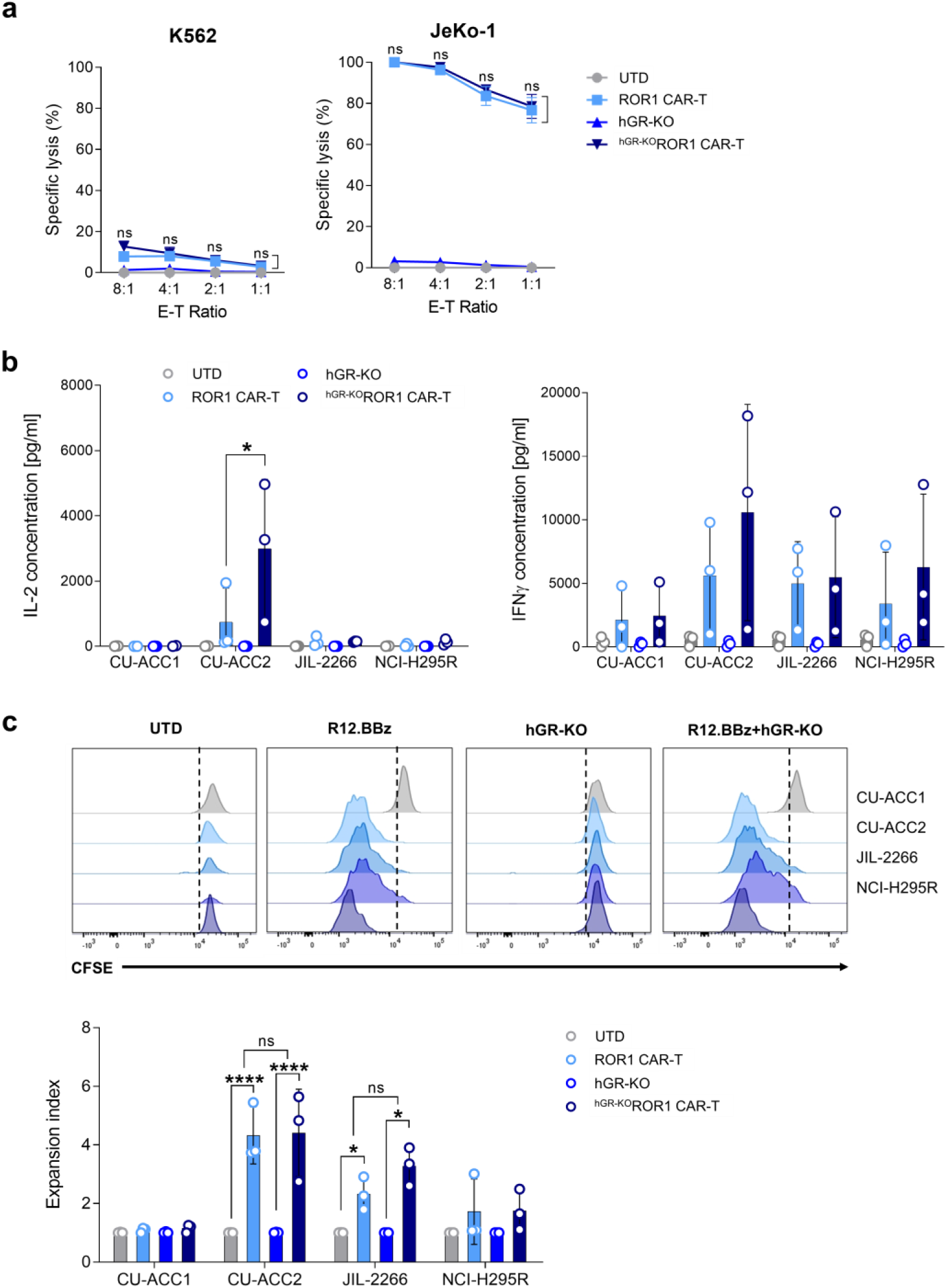
Anti-tumour reactivity of GC-resistant ROR1 CAR-T cells *in vitro*. (**a**) Specific lysis of ROR1^+^ JeKo-1 mantle cell lymphoma and ROR1^−^ K562 leukemia cell line with different effector-to-target ratios (E-T) after 24 hours. (**b**) IL-2 and IFN-γ cytokine secretion of ROR1 CAR-T cells with and without hGR knockout after antigen contact for 24 hours was assessed by ELISA in the supernatant. (**c**) Proliferation and expansion of ROR1 CAR-T cells upon antigen contact in all four ACC cell lines after 72 hours assessed by FACS-analysis. Statistical analyses were considered significant if p-value was *p<0.05,**p<0.01,***p<0.001, ****p<0.0001; ns, not significant; hGR, human glucocorticoid receptor; ELISA, enzyme-linked immunosorbent assay; FACS, fluorescence activated cell sorting.

**Extended Data Fig. 9.**
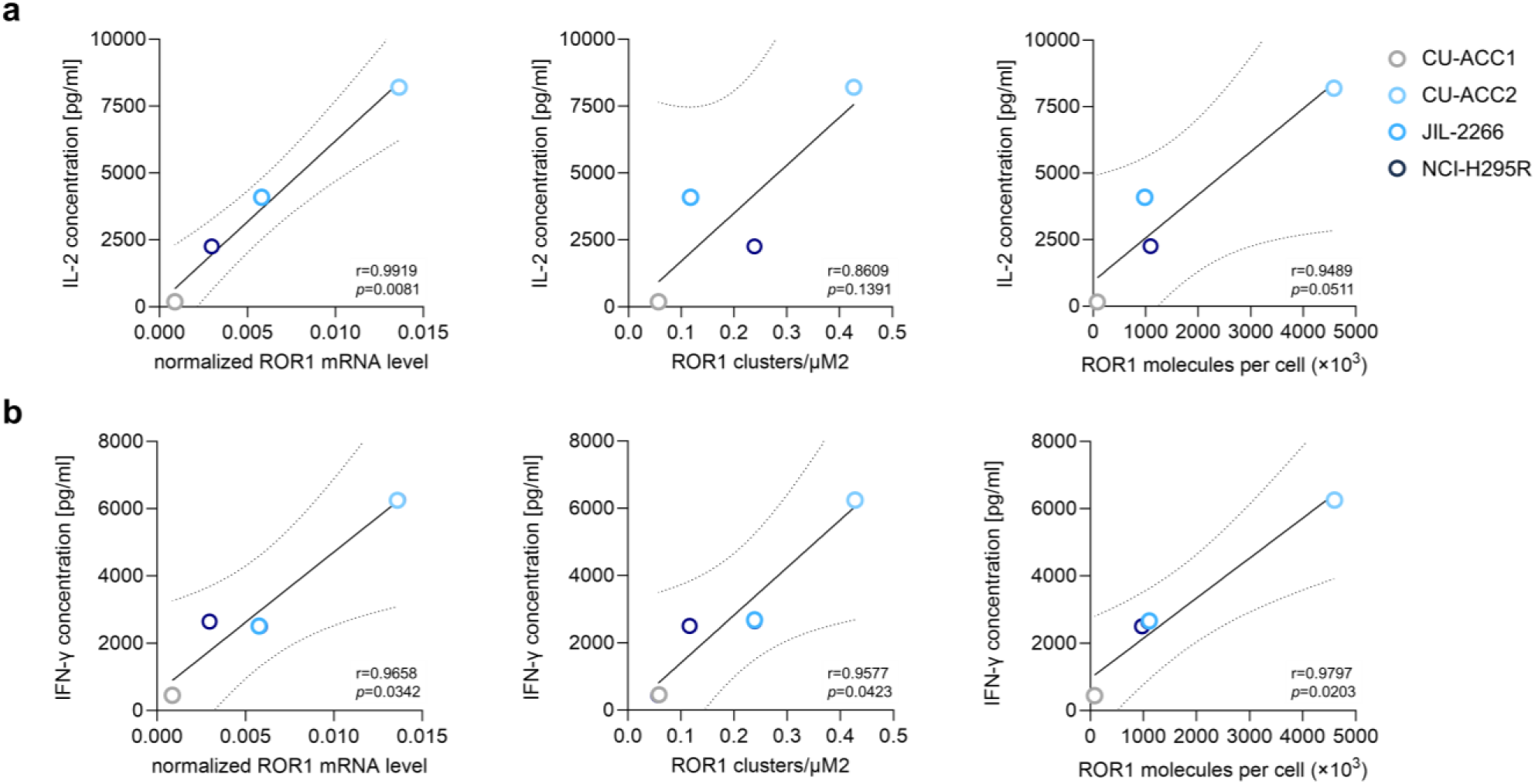
Correlation between ROR1 antigen density and cytokine production from ^hGR-KO^ROR1 CAR-T cells. (**a**) IL-2 and (**b**) IFN-γ cytokine secretion of ^hGR-KO^ROR1 CAR-T cells correlate with ROR1 antigen levels (RNA, density and molecule count) in ACC cell lines.

**Extended Data Fig. 10.**
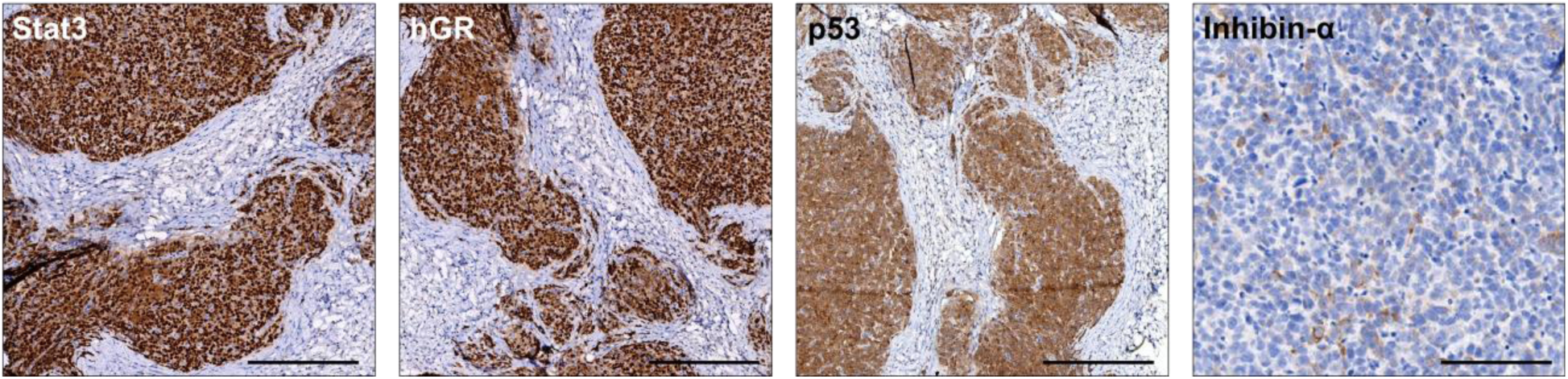
Phenotype of ACC xenografts in the NSG mouse model. Immunohistochemical staining of Stat3, hGR, p53 and inhibin-α on ACC tumours from the steroidogenic ACC mouse model (NSG/NCI-H295R).

**Extended Data Fig. 11.**
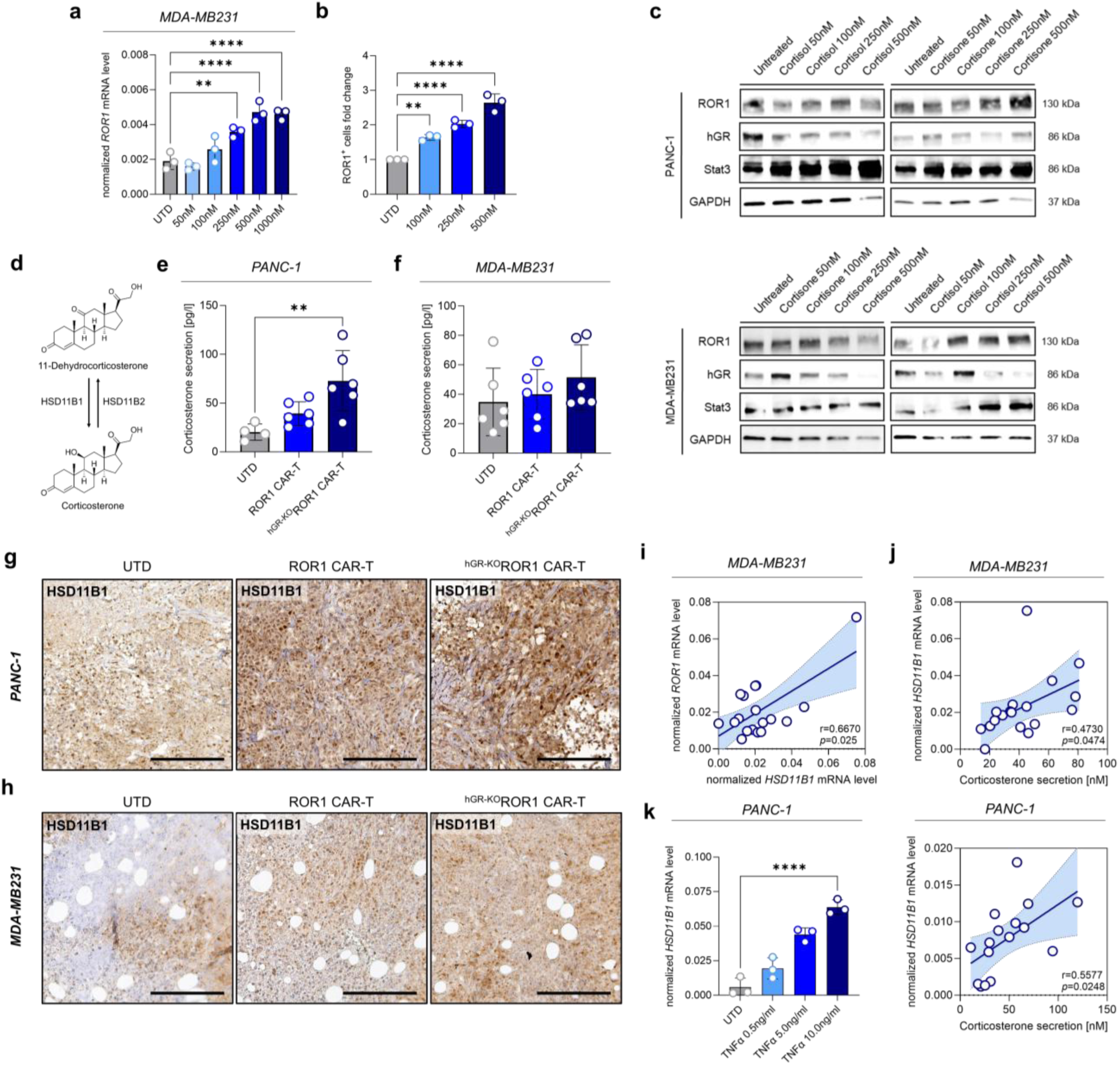
^hGR-KO^ROR1 CAR-T cells are effective against TNBC and PDAC *in vivo*. (**a**) ROR1 mRNA expression after different doses of cortisone (UTD, untreated control). (**b**) Conversion from cortisone-into-cortisol upregulates ROR1 protein cell surface expression assessed by FACS analysis. (**c**) ROR1. hGR and STAT3 protein expression after increasing doses of cortisol and cortisone in PANC-1 and MDA-MB231 cells. (**d**) Scheme that demonstrates the molecular recouperation from 11-dehydrocorticosterone-into-corticosterone through HSD11B1. (**e** and **f**) Serum corticosterone levels in PDAC and TNBC bearing mice in all three different treatment groups. (**g**) and (**h**) show representative immunohistochemistry pictures of ROR1 and HSD11B1 protein expression at *in vivo* endpoint analyses from PDAC and TNBC tumours. (**i**) Association between ROR1 and HSD11B1 mRNA expression in TNBC tumours. (**j**) Association between HSD11B1 mRNA expression and serum corticosterone levels in mice. (**k**) HSD11B1 mRNA expression in PANC-1 cells after increasing doses of TNFα. Statistical analyses were performed using one-way ANOVA with Dunnett’s test for multiple comparisons (**a,b,c,e,j**) and Pearson correlation (**h,i**) and were considered significant if *p*-value was \**p*<0.05,\*\**p*<0.01,\*\*\**p*<0.001, \*\*\*\**p*<0.0001; ns, not significant.

**Extended Data Fig. 12.**
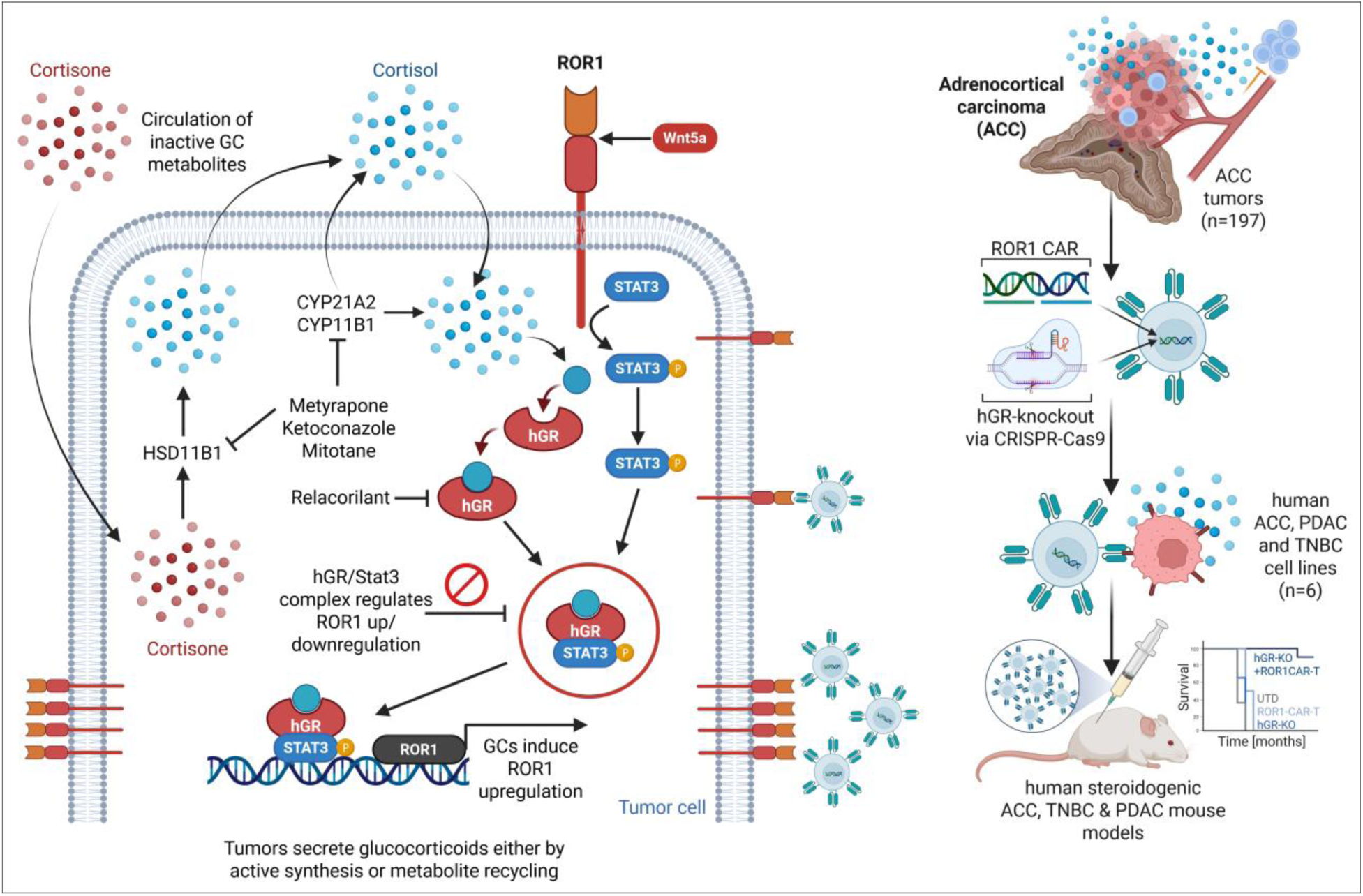
Graphical abstract. Glucocorticoid excess in endocrine and non-endocrine cancers fuels an autocrine signaling loop that augments expression of the oncogenic driver ROR1 for targeting with ROR1 CAR-T cells that are shielded from the paracrine immunosuppressive effects of cancer-derived GCs through hGR knockout.

## REFERENCES AND NOTES

1. Santomasso, B.D., Nastoupil, L.J., Adkins, S., Lacchetti, C., Schneider, B.J., Anadkat, M., Atkins, M.B., Brassil, K.J., Caterino, J.M., Chau, I., et al. (2021). Management of Immune-Related Adverse Events in Patients Treated With Chimeric Antigen Receptor T-Cell Therapy: ASCO Guideline. J Clin Oncol 39, 3978–3992. 10.1200/JCO.21.01992.

2. Schneider, B.J., Naidoo, J., Santomasso, B.D., Lacchetti, C., Adkins, S., Anadkat, M., Atkins, M.B., Brassil, K.J., Caterino, J.M., Chau, I., et al. (2021). Management of Immune-Related Adverse Events in Patients Treated With Immune Checkpoint Inhibitor Therapy: ASCO Guideline Update. J Clin Oncol 39, 4073–4126. 10.1200/JCO.21.01440.

3. Cain, D.W., and Cidlowski, J.A. (2017). Immune regulation by glucocorticoids. Nat Rev Immunol 17, 233–247. 10.1038/nri.2017.1.

4. Taves, M.D., and Ashwell, J.D. (2021). Glucocorticoids in T cell development, differentiation and function. Nat Rev Immunol 21, 233–243. 10.1038/s41577-020-00464-0.

5. Arbour, K.C., Mezquita, L., Long, N., Rizvi, H., Auclin, E., Ni, A., Martinez-Bernal, G., Ferrara, R., Lai, W.V., Hendriks, L.E.L., et al. (2018). Impact of Baseline Steroids on Efficacy of Programmed Cell Death-1 and Programmed Death-Ligand 1 Blockade in Patients With Non-Small-Cell Lung Cancer. J Clin Oncol 36, 2872–2878. 10.1200/JCO.2018.79.0006.

6. Strati, P., Ahmed, S., Furqan, F., Fayad, L.E., Lee, H.J., Iyer, S.P., Nair, R., Nastoupil, L.J., Parmar, S., Rodriguez, M.A., et al. (2021). Prognostic impact of corticosteroids on efficacy of chimeric antigen receptor T-cell therapy in large B-cell lymphoma. Blood 137, 3272–3276. 10.1182/blood.2020008865.

7. Yang, H., Xia, L., Chen, J., Zhang, S., Martin, V., Li, Q., Lin, S., Chen, J., Calmette, J., Lu, M., et al. (2019). Stress-glucocorticoid-TSC22D3 axis compromises therapy-induced antitumor immunity. Nat Med 25, 1428–1441. 10.1038/s41591-019-0566-4.

8. Acharya, N., Madi, A., Zhang, H., Klapholz, M., Escobar, G., Dulberg, S., Christian, E., Ferreira, M., Dixon, K.O., Fell, G., et al. (2020). Endogenous Glucocorticoid Signaling Regulates CD8(+) T Cell Differentiation and Development of Dysfunction in the Tumor Microenvironment. Immunity 53, 658–671.e656. 10.1016/j.immuni.2020.08.005.

9. Taves, M.D., and Ashwell, J.D. (2021). Glucocorticoids in T cell development, differentiation and function. Nature Reviews Immunology 21, 233–243. 10.1038/s41577-020-00464-0.

10. Swatler, J., Ju, Y.J., Anderson, A.C., and Lugli, E. (2023). Tumors recycle glucocorticoids to drive Treg-mediated immunosuppression. J Clin Invest 133. 10.1172/JCI173141.

11. Taves, M.D., Otsuka, S., Taylor, M.A., Donahue, K.M., Meyer, T.J., Cam, M.C., and Ashwell, J.D. (2023). Tumors produce glucocorticoids by metabolite recycling, not synthesis, and activate Tregs to promote growth. The Journal of Clinical Investigation 133. 10.1172/JCI164599.

12. Fassnacht, M., Assie, G., Baudin, E., Eisenhofer, G., de la Fouchardiere, C., Haak, H.R., de Krijger, R., Porpiglia, F., Terzolo, M., Berruti, A., and, E.G.C.E.a. (2020). Adrenocortical carcinomas and malignant phaeochromocytomas: ESMO-EURACAN Clinical Practice Guidelines for diagnosis, treatment and follow-up. Ann Oncol 31, 1476–1490. 10.1016/j.annonc.2020.08.2099.

13. Landwehr, L.S., Altieri, B., Sbiera, I., Remde, H., Kircher, S., Olabe, J., Sbiera, S., Kroiss, M., and Fassnacht, M. (2024). Expression and Prognostic Relevance of PD-1, PD-L1, and CTLA-4 Immune Checkpoints in Adrenocortical Carcinoma. J Clin Endocrinol Metab 109, 2325–2334. 10.1210/clinem/dgae109.

14. Landwehr, L.S., Altieri, B., Schreiner, J., Sbiera, I., Weigand, I., Kroiss, M., Fassnacht, M., and Sbiera, S. (2020). Interplay between glucocorticoids and tumor-infiltrating lymphocytes on the prognosis of adrenocortical carcinoma. J Immunother Cancer 8. 10.1136/jitc-2019-000469.

15. Martins Nascentes Melo, L., Herrera-Rios, D., Hinze, D., Loffek, S., Oezel, I., Turiello, R., Klein, J., Leonardelli, S., Westedt, I.V., Al-Matary, Y., et al. (2023). Glucocorticoid activation by HSD11B1 limits T cell-driven interferon signaling and response to PD-1 blockade in melanoma. J Immunother Cancer 11. 10.1136/jitc-2021-004150.

16. Zhang, H., Liu, A., Bo, W., Zhang, M., Wang, H., Feng, X., and Wu, Y. (2024). Upregulation of HSD11B1 promotes cortisol production and inhibits NK cell activation in pancreatic adenocarcinoma. Mol Immunol 175, 10–19. 10.1016/j.molimm.2024.08.005.

17. Timperi, E., Gueguen, P., Molgora, M., Magagna, I., Kieffer, Y., Lopez-Lastra, S., Sirven, P., Baudrin, L.G., Baulande, S., Nicolas, A., et al. (2022). Lipid-Associated Macrophages Are Induced by Cancer-Associated Fibroblasts and Mediate Immune Suppression in Breast Cancer. Cancer Res 82, 3291–3306. 10.1158/0008-5472.CAN-22-1427.

18. Mahata, B., Pramanik, J., van der Weyden, L., Polanski, K., Kar, G., Riedel, A., Chen, X., Fonseca, N.A., Kundu, K., Campos, L.S., et al. (2020). Tumors induce de novo steroid biosynthesis in T cells to evade immunity. Nat Commun 11, 3588. 10.1038/s41467-020-17339-6.

19. Taves, M.D., Otsuka, S., Taylor, M.A., Donahue, K.M., Meyer, T.J., Cam, M.C., and Ashwell, J.D. (2023). Tumors produce glucocorticoids by metabolite recycling, not synthesis, and activate Tregs to promote growth. J Clin Invest 133. 10.1172/JCI164599.

20. Labanieh, L., and Mackall, C.L. (2023). CAR immune cells: design principles, resistance and the next generation. Nature 614, 635–648. 10.1038/s41586-023-05707-3.

21. Giordano, T.J., Kuick, R., Else, T., Gauger, P.G., Vinco, M., Bauersfeld, J., Sanders, D., Thomas, D.G., Doherty, G., and Hammer, G. (2009). Molecular classification and prognostication of adrenocortical tumors by transcriptome profiling. Clin Cancer Res 15, 668–676. 10.1158/1078-0432.CCR-08-1067.

22. Fassnacht, M., Johanssen, S., Quinkler, M., Bucsky, P., Willenberg, H.S., Beuschlein, F., Terzolo, M., Mueller, H.H., Hahner, S., Allolio, B., et al. (2009). Limited prognostic value of the 2004 International Union Against Cancer staging classification for adrenocortical carcinoma: proposal for a Revised TNM Classification. Cancer 115, 243–250. 10.1002/cncr.24030.

23. Fassnacht, M., Terzolo, M., Allolio, B., Baudin, E., Haak, H., Berruti, A., Welin, S., Schade-Brittinger, C., Lacroix, A., Jarzab, B., et al. (2012). Combination chemotherapy in advanced adrenocortical carcinoma. N Engl J Med 366, 2189–2197. 10.1056/NEJMoa1200966.

24. Fassnacht, M., Assie, G., Baudin, E., Eisenhofer, G., de la Fouchardiere, C., Haak, H.R., de Krijger, R., Porpiglia, F., Terzolo, M., and Berruti, A. (2020). Adrenocortical carcinomas and malignant phaeochromocytomas: ESMO-EURACAN Clinical Practice Guidelines for diagnosis, treatment and follow-up. Annals of Oncology 31, 1476–1490. 10.1016/j.annonc.2020.08.2099.

25. Kiseljak-Vassiliades, K., Zhang, Y., Bagby, S.M., Kar, A., Pozdeyev, N., Xu, M., Gowan, K., Sharma, V., Raeburn, C.D., Albuja-Cruz, M., et al. (2018). Development of new preclinical models to advance adrenocortical carcinoma research. Endocr Relat Cancer 25, 437–451. 10.1530/ERC-17-0447.

26. Rainey, W.E., Saner, K., and Schimmer, B.P. (2004). Adrenocortical cell lines. Mol Cell Endocrinol 228, 23–38. 10.1016/j.mce.2003.12.020.

27. Landwehr, L.S., Schreiner, J., Appenzeller, S., Kircher, S., Herterich, S., Sbiera, S., Fassnacht, M., Kroiss, M., and Weigand, I. (2021). A novel patient-derived cell line of adrenocortical carcinoma shows a pathogenic role of germline MUTYH mutation and high tumour mutational burden. Eur J Endocrinol 184, 823–835. 10.1530/EJE-20-1423.

28. Conway, M.E., McDaniel, J.M., Graham, J.M., Guillen, K.P., Oliver, P.G., Parker, S.L., Yue, P., Turkson, J., Buchsbaum, D.J., Welm, B.E., et al. (2020). STAT3 and GR Cooperate to Drive Gene Expression and Growth of Basal-Like Triple-Negative Breast Cancer. Cancer Res 80, 4355–4370. 10.1158/0008-5472.CAN-20-1379.

29. Zhang, S., Chen, L., Cui, B., Chuang, H.-Y., Yu, J., Wang-Rodriguez, J., Tang, L., Chen, G., Basak, G.W., and Kipps, T.J. (2012). ROR1 Is Expressed in Human Breast Cancer and Associated with Enhanced Tumor-Cell Growth. PLoS One 7, e31127. 10.1371/journal.pone.0031127.

30. Liu, D., Sharbeen, G., Phillips, P., and Ford, C.E. (2021). ROR1 and ROR2 expression in pancreatic cancer. BMC Cancer 21, 1199. 10.1186/s12885-021-08952-9.

31. Sampath-Kumar, R., Yu, M., Khalil, M.W., and Yang, K. (1997). Metyrapone is a competitive inhibitor of 11beta-hydroxysteroid dehydrogenase type 1 reductase. J Steroid Biochem Mol Biol 62, 195–199. 10.1016/s0960-0760(97)00027-7.

32. Da Via, M.C., Dietrich, O., Truger, M., Arampatzi, P., Duell, J., Heidemeier, A., Zhou, X., Danhof, S., Kraus, S., Chatterjee, M., et al. (2021). Homozygous BCMA gene deletion in response to anti-BCMA CAR T cells in a patient with multiple myeloma. Nat Med 27, 616–619. 10.1038/s41591-021-01245-5.

33. Spiegel, J.Y., Patel, S., Muffly, L., Hossain, N.M., Oak, J., Baird, J.H., Frank, M.J., Shiraz, P., Sahaf, B., Craig, J., et al. (2021). CAR T cells with dual targeting of CD19 and CD22 in adult patients with recurrent or refractory B cell malignancies: a phase 1 trial. Nat Med 27, 1419–1431. 10.1038/s41591-021-01436-0.

34. Von Knebel Doeberitz, M., Koch, S., Drzonek, H., and Zur Hausen, H. (1990). Glucocorticoid hormones reduce the expression of major histocompatibility class I antigens on human epithelial cells. Eur J Immunol 20, 35–40. 10.1002/eji.1830200106.

35. Galon, J., Franchimont, D., Hiroi, N., Frey, G., Boettner, A., Ehrhart-Bornstein, M., O’Shea, J.J., Chrousos, G.P., and Bornstein, S.R. (2002). Gene profiling reveals unknown enhancing and suppressive actions of glucocorticoids on immune cells. FASEB J 16, 61–71. 10.1096/fj.01-0245com.

36. Zhai, E., Liang, W., Lin, Y., Huang, L., He, X., Cai, S., Chen, J., Zhang, N., Li, J., Zhang, Q., et al. (2018). HSP70/HSP90-Organizing Protein Contributes to Gastric Cancer Progression in an Autocrine Fashion and Predicts Poor Survival in Gastric Cancer. Cell Physiol Biochem 47, 879–892. 10.1159/000490080.

37. Lin, Y., Zhai, E., Liao, B., Xu, L., Zhang, X., Peng, S., He, Y., Cai, S., Zeng, Z., and Chen, M. (2017). Autocrine VEGF signaling promotes cell proliferation through a PLC-dependent pathway and modulates Apatinib treatment efficacy in gastric cancer. Oncotarget 8, 11990–12002. 10.18632/oncotarget.14467.

38. He, Z., Song, B., Zhu, M., and Liu, J. (2023). Comprehensive pan-cancer analysis of STAT3 as a prognostic and immunological biomarker. Sci Rep 13, 5069. 10.1038/s41598-023-31226-2.

39. Menck, K., Heinrichs, S., Baden, C., and Bleckmann, A. (2021). The WNT/ROR Pathway in Cancer: From Signaling to Therapeutic Intervention. Cells 10. 10.3390/cells10010142.

40. Cui, B., Zhang, S., Chen, L., Yu, J., Widhopf, G.F., 2nd, Fecteau, J.F., Rassenti, L.Z., and Kipps, T.J. (2013). Targeting ROR1 inhibits epithelial-mesenchymal transition and metastasis. Cancer Res 73, 3649–3660. 10.1158/0008-5472.CAN-12-3832.

41. Balakrishnan, A., Goodpaster, T., Randolph-Habecker, J., Hoffstrom, B.G., Jalikis, F.G., Koch, L.K., Berger, C., Kosasih, P.L., Rajan, A., Sommermeyer, D., et al. (2017). Analysis of ROR1 Protein Expression in Human Cancer and Normal Tissues. Clin Cancer Res 23, 3061–3071. 10.1158/1078-0432.CCR-16-2083.

42. Zhang, S., Chen, L., Wang-Rodriguez, J., Zhang, L., Cui, B., Frankel, W., Wu, R., and Kipps, T.J. (2012). The onco-embryonic antigen ROR1 is expressed by a variety of human cancers. Am J Pathol 181, 1903–1910. 10.1016/j.ajpath.2012.08.024.

43. Hudecek, M., Lupo-Stanghellini, M.-T., Kosasih, P.L., Sommermeyer, D., Jensen, M.C., Rader, C., and Riddell, S.R. (2013). Receptor Affinity and Extracellular Domain Modifications Affect Tumor Recognition by ROR1-Specific Chimeric Antigen Receptor T Cells. Clinical Cancer Research 19, 3153–3164. 10.1158/1078-0432.Ccr-13-0330.

44. Wallstabe, L., Gottlich, C., Nelke, L.C., Kuhnemundt, J., Schwarz, T., Nerreter, T., Einsele, H., Walles, H., Dandekar, G., Nietzer, S.L., and Hudecek, M. (2019). ROR1-CAR T cells are effective against lung and breast cancer in advanced microphysiologic 3D tumor models. JCI Insight 4. 10.1172/jci.insight.126345.

45. Obradovic, M.M.S., Hamelin, B., Manevski, N., Couto, J.P., Sethi, A., Coissieux, M.M., Munst, S., Okamoto, R., Kohler, H., Schmidt, A., and Bentires-Alj, M. (2019). Glucocorticoids promote breast cancer metastasis. Nature 567, 540–544. 10.1038/s41586-019-1019-4.

46. Karvonen, H., Arjama, M., Kaleva, L., Niininen, W., Barker, H., Koivisto-Korander, R., Tapper, J., Pakarinen, P., Lassus, H., Loukovaara, M., et al. (2020). Glucocorticoids induce differentiation and chemoresistance in ovarian cancer by promoting ROR1-mediated stemness. Cell Death Dis 11, 790. 10.1038/s41419-020-03009-4.

47. Yu, J., Chen, L., Cui, B., Widhopf, G.F., II, Shen, Z., Wu, R., Zhang, L., Zhang, S., Briggs, S.P., and Kipps, T.J. (2016). Wnt5a induces ROR1/ROR2 heterooligomerization to enhance leukemia chemotaxis and proliferation. The Journal of Clinical Investigation 126, 585–598. 10.1172/JCI83535.

48. Rozovski, U., Harris, D.M., Li, P., Liu, Z., Jain, P., Ferrajoli, A., Burger, J.A., Bose, P., Thompson, P.A., Jain, N., et al. (2019). STAT3-Induced Wnt5a Provides Chronic Lymphocytic Leukemia Cells with Survival Advantage. J Immunol 203, 3078–3085. 10.4049/jimmunol.1900389.

49. Yu, J., Chen, L., Cui, B., Wu, C., Choi, M.Y., Chen, Y., Zhang, L., Rassenti, L.Z., Widhopf Ii, G.F., and Kipps, T.J. (2017). Cirmtuzumab inhibits Wnt5a-induced Rac1 activation in chronic lymphocytic leukemia treated with ibrutinib. Leukemia 31, 1333–1339. 10.1038/leu.2016.368.

50. Thorsson, V., Gibbs, D.L., Brown, S.D., Wolf, D., Bortone, D.S., Ou Yang, T.H., Porta-Pardo, E., Gao, G.F., Plaisier, C.L., Eddy, J.A., et al. (2018). The Immune Landscape of Cancer. Immunity 48, 812–830.e814. 10.1016/j.immuni.2018.03.023.

51. Brummer, A.B., Yang, X., Ma, E., Gutova, M., Brown, C.E., and Rockne, R.C. (2022). Dose-dependent thresholds of dexamethasone destabilize CAR T-cell treatment efficacy. PLoS Comput Biol 18, e1009504. 10.1371/journal.pcbi.1009504.

52. Poiret, T., Vikberg, S., Schoutrop, E., Mattsson, J., and Magalhaes, I. (2024). CAR T cells and T cells phenotype and function are impacted by glucocorticoid exposure with different magnitude. Journal of Translational Medicine 22, 273. 10.1186/s12967-024-05063-4.

53. Santomasso, B.D., Nastoupil, L.J., Adkins, S., Lacchetti, C., Schneider, B.J., Anadkat, M., Atkins, M.B., Brassil, K.J., Caterino, J.M., Chau, I., et al. (2021). Management of Immune-Related Adverse Events in Patients Treated With Chimeric Antigen Receptor T-Cell Therapy: ASCO Guideline. Journal of Clinical Oncology 39, 3978–3992. 10.1200/jco.21.01992.

54. Hayden, P.J., Roddie, C., Bader, P., Basak, G.W., Bonig, H., Bonini, C., Chabannon, C., Ciceri, F., Corbacioglu, S., Ellard, R., et al. (2022). Management of adults and children receiving CAR T-cell therapy: 2021 best practice recommendations of the European Society for Blood and Marrow Transplantation (EBMT) and the Joint Accreditation Committee of ISCT and EBMT (JACIE) and the European Haematology Association (EHA). Ann Oncol 33, 259–275. 10.1016/j.annonc.2021.12.003.

55. Giavridis, T., van der Stegen, S.J.C., Eyquem, J., Hamieh, M., Piersigilli, A., and Sadelain, M. (2018). CAR T cell-induced cytokine release syndrome is mediated by macrophages and abated by IL-1 blockade. Nat Med 24, 731–738. 10.1038/s41591-018-0041-7.

56. Norelli, M., Camisa, B., Barbiera, G., Falcone, L., Purevdorj, A., Genua, M., Sanvito, F., Ponzoni, M., Doglioni, C., Cristofori, P., et al. (2018). Monocyte-derived IL-1 and IL-6 are differentially required for cytokine-release syndrome and neurotoxicity due to CAR T cells. Nat Med 24, 739–748. 10.1038/s41591-018-0036-4.

57. Gust, J., Hay, K.A., Hanafi, L.A., Li, D., Myerson, D., Gonzalez-Cuyar, L.F., Yeung, C., Liles, W.C., Wurfel, M., Lopez, J.A., et al. (2017). Endothelial Activation and Blood-Brain Barrier Disruption in Neurotoxicity after Adoptive Immunotherapy with CD19 CAR-T Cells. Cancer Discov 7, 1404–1419. 10.1158/2159-8290.Cd-17-0698.

58. Mazein, A., Lopata, O., Reiche, K., Sewald, K., Alb, M., Sakellariou, C., Gogesch, P., Morgan, H., Neuhaus, V., Pham, N.N., et al. (2025). An explorable model of an adverse outcome pathway of cytokine release syndrome related to the administration of immunomodulatory biotherapeutics and cellular therapies. Front Immunol 16, 1601670. 10.3389/fimmu.2025.1601670.

59. Kaeuferle, T., Deisenberger, L., Jablonowski, L., Stief, T.A., Blaeschke, F., Willier, S., and Feuchtinger, T. (2020). CRISPR-Cas9-Mediated Glucocorticoid Resistance in Virus-Specific T Cells for Adoptive T Cell Therapy Posttransplantation. Mol Ther 28, 1965–1973. 10.1016/j.ymthe.2020.06.002.

60. Brown, C.E., Rodriguez, A., Palmer, J., Ostberg, J.R., Naranjo, A., Wagner, J.R., Aguilar, B., Starr, R., Weng, L., Synold, T.W., et al. (2022). Off-the-shelf, steroid-resistant, IL13Rα2-specific CAR T cells for treatment of glioblastoma. Neuro Oncol 24, 1318–1330. 10.1093/neuonc/noac024.

61. Paszkiewicz, P.J., Frassle, S.P., Srivastava, S., Sommermeyer, D., Hudecek, M., Drexler, I., Sadelain, M., Liu, L., Jensen, M.C., Riddell, S.R., and Busch, D.H. (2016). Targeted antibody-mediated depletion of murine CD19 CAR T cells permanently reverses B cell aplasia. J Clin Invest 126, 4262–4272. 10.1172/JCI84813.

62. Mestermann, K., Giavridis, T., Weber, J., Rydzek, J., Frenz, S., Nerreter, T., Mades, A., Sadelain, M., Einsele, H., and Hudecek, M. (2019). The tyrosine kinase inhibitor dasatinib acts as a pharmacologic on/off switch for CAR T cells. Sci Transl Med 11. 10.1126/scitranslmed.aau5907.

63. Taves, M.D., Gomez-Sanchez, C.E., and Soma, K.K. (2011). Extra-adrenal glucocorticoids and mineralocorticoids: evidence for local synthesis, regulation, and function. Am J Physiol Endocrinol Metab 301, E11–24. 10.1152/ajpendo.00100.2011.

64. Deng, Y., Xia, X., Zhao, Y., Zhao, Z., Martinez, C., Yin, W., Yao, J., Hang, Q., Wu, W., Zhang, J., et al. (2021). Glucocorticoid receptor regulates PD-L1 and MHC-I in pancreatic cancer cells to promote immune evasion and immunotherapy resistance. Nature Communications 12, 7041. 10.1038/s41467-021-27349-7.

65. Sidler, D., Renzulli, P., Schnoz, C., Berger, B., Schneider-Jakob, S., Fluck, C., Inderbitzin, D., Corazza, N., Candinas, D., and Brunner, T. (2011). Colon cancer cells produce immunoregulatory glucocorticoids. Oncogene 30, 2411–2419. 10.1038/onc.2010.629.

66. Ahmed, A., Reinhold, C., Breunig, E., Phan, T.S., Dietrich, L., Kostadinova, F., Urwyler, C., Merk, V.M., Noti, M., Toja da Silva, I., et al. (2023). Immune escape of colorectal tumours via local LRH-1/Cyp11b1-mediated synthesis of immunosuppressive glucocorticoids. Mol Oncol 17, 1545–1566. 10.1002/1878-0261.13414.

67. Fassnacht, M., Johanssen, S., Quinkler, M., Bucsky, P., Willenberg, H.S., Beuschlein, F., Terzolo, M., Mueller, H.H., Hahner, S., Allolio, B., et al. (2009). Limited prognostic value of the 2004 International Union Against Cancer staging classification for adrenocortical carcinoma: proposal for a Revised TNM Classification. Cancer 115, 243–250. 10.1002/cncr.24030.

68. Sbiera, I., Kircher, S., Altieri, B., Lenz, K., Hantel, C., Fassnacht, M., Sbiera, S., and Kroiss, M. (2021). Role of FGF Receptors and Their Pathways in Adrenocortical Tumors and Possible Therapeutic Implications. Front Endocrinol (Lausanne) 12, 795116. 10.3389/fendo.2021.795116.

69. Gomez-Sanchez, C.E., Qi, X., Velarde-Miranda, C., Plonczynski, M.W., Parker, C.R., Rainey, W., Satoh, F., Maekawa, T., Nakamura, Y., Sasano, H., and Gomez-Sanchez, E.P. (2014). Development of monoclonal antibodies against human CYP11B1 and CYP11B2. Mol Cell Endocrinol 383, 111–117. 10.1016/j.mce.2013.11.022.

70. Wolter, S., Loschberger, A., Holm, T., Aufmkolk, S., Dabauvalle, M.C., van de Linde, S., and Sauer, M. (2012). rapidSTORM: accurate, fast open-source software for localization microscopy. Nat Methods 9, 1040–1041. 10.1038/nmeth.2224.

71. Doose, S. (2022). LOCAN: a python library for analyzing single-molecule localization microscopy data. Bioinformatics 38, 2670–2672. 10.1093/bioinformatics/btac160.

72. Kroiss, M., Reuss, M., Kuhner, D., Johanssen, S., Beyer, M., Zink, M., Hartmann, M.F., Dhir, V., Wudy, S.A., Arlt, W., et al. (2011). Sunitinib Inhibits Cell Proliferation and Alters Steroidogenesis by Down-Regulation of HSD3B2 in Adrenocortical Carcinoma Cells. Front Endocrinol (Lausanne) 2, 27. 10.3389/fendo.2011.00027.

73. Altieri, B., Sbiera, S., Della Casa, S., Weigand, I., Wild, V., Steinhauer, S., Fadda, G., Kocot, A., Bekteshi, M., Mambretti, E.M., et al. (2017). Livin/BIRC7 expression as malignancy marker in adrenocortical tumors. Oncotarget 8, 9323–9338. 10.18632/oncotarget.14067.

74. Hudecek, M., Schmitt, T.M., Baskar, S., Lupo-Stanghellini, M.T., Nishida, T., Yamamoto, T.N., Bleakley, M., Turtle, C.J., Chang, W.C., Greisman, H.A., et al. (2010). The B-cell tumor-associated antigen ROR1 can be targeted with T cells modified to express a ROR1-specific chimeric antigen receptor. Blood 116, 4532–4541. 10.1182/blood-2010-05-283309.

75. Hudecek, M., Sommermeyer, D., Kosasih, P.L., Silva-Benedict, A., Liu, L., Rader, C., Jensen, M.C., and Riddell, S.R. (2015). The nonsignaling extracellular spacer domain of chimeric antigen receptors is decisive for in vivo antitumor activity. Cancer Immunol Res 3, 125–135. 10.1158/2326-6066.CIR-14-0127.

76. Kang, J., Lee, J.H., Cha, H., An, J., Kwon, J., Lee, S., Kim, S., Baykan, M.Y., Kim, S.Y., An, D., et al. (2024). Systematic dissection of tumor-normal single-cell ecosystems across a thousand tumors of 30 cancer types. Nat Commun 15, 4067. 10.1038/s41467-024-48310-4.

77. Bilous, M., Tran, L., Cianciaruso, C., Gabriel, A., Michel, H., Carmona, S.J., Pittet, M.J., and Gfeller, D. (2022). Metacells untangle large and complex single-cell transcriptome networks. BMC Bioinformatics 23, 336. 10.1186/s12859-022-04861-1.

78. Hao, Y., Hao, S., Andersen-Nissen, E., Mauck, W.M., 3rd, Zheng, S., Butler, A., Lee, M.J., Wilk, A.J., Darby, C., Zager, M., et al. (2021). Integrated analysis of multimodal single-cell data. Cell 184, 3573–3587 e3529. 10.1016/j.cell.2021.04.048.

79. Korsunsky, I., Millard, N., Fan, J., Slowikowski, K., Zhang, F., Wei, K., Baglaenko, Y., Brenner, M., Loh, P.R., and Raychaudhuri, S. (2019). Fast, sensitive and accurate integration of single-cell data with Harmony. Nat Methods 16, 1289–1296. 10.1038/s41592-019-0619-0.

